# Myoinhibitory peptide regulates feeding in the marine annelid *Platynereis*

**DOI:** 10.1101/008854

**Authors:** Elizabeth A. Williams, Markus Conzelmann, Gáspár Jékely

## Abstract

**Background:** During larval settlement and metamorphosis, marine invertebrates undergo changes in habitat, morphology, behavior and physiology. This change between life-cycle stages is often associated with a change in diet or a transition between a non-feeding and a feeding form. How larvae regulate changes in feeding during this life cycle transition is not well understood. Neuropeptides are known to regulate several aspects of feeding, such as food search, ingestion and digestion. The marine annelid *Platynereis dumerilii* has a complex life cycle with a pelagic non-feeding larval stage and a benthic feeding postlarval stage, linked by the process of settlement. The conserved neuropeptide myoinhibitory peptide (MIP) is a key regulator of larval settlement behavior in *Platynereis*. Whether MIP also regulates the initiation of feeding, another aspect of the pelagic-to-benthic transition in *Platynereis*, is currently unknown.

**Results:** Here, we explore the contribution of MIP to feeding in settled *Platynereis* postlarvae. We find that MIP is expressed in the gut of developing larvae in sensory neurons that densely innervate the foregut and hindgut. Activating MIP signaling by synthetic neuropeptide addition causes increased gut peristalsis and more frequent pharynx extensions leading to increased food intake. Conversely, morpholino-mediated knockdown of MIP expression inhibits feeding. In the long-term, treatment of *Platynereis* postlarvae with synthetic MIP increases growth rate and results in earlier cephalic metamorphosis.

**Conclusions:** Our results show that MIP activates ingestion and digestion in *Platynereis* postlarvae. MIP is expressed in sensory-neurosecretory cells of the digestive system indicating that following larval settlement, feeding is initiated by a direct sensory mechanism. This is similar to the mechanism by which MIP induces larval settlement. The pleiotropic roles of MIP may thus have evolved by redeploying the same signaling mechanism in different aspects of a life cycle transition.

## Background

Many organisms have a complex life cycle consisting of distinct stages that differ in form, physiology, behavior and habitat. Among benthic marine invertebrates, a common life cycle strategy, the biphasic life cycle, consists of a free-swimming larva that settles to the ocean floor and undergoes metamorphosis to a bottom-dwelling adult [1]. Marine invertebrate larval settlement is often coupled to the initiation of feeding or a change in diet [2]. These behavioral, physiological and morphological changes have to be tightly coordinated for the successful transition to a benthic life style. Knowledge of how this transition is regulated is important for understanding population structure in the ocean, life-history evolution, and how the environment influences animal life cycles [3–8].

The marine annelid *Platynereis dumerilii* is a powerful marine invertebrate model for studying the molecular details of marine larval behavior, including settlement [9–12]. *Platynereis* has a biphasic life cycle with free-swimming, non-feeding larval (trochophore and nectochaete) stages and bottom-dwelling feeding postlarval, juvenile and adult stages [13]. Larval settlement is followed by a period of growth and feeding, during which the juvenile *Platynereis* add additional posterior segments. Cephalic metamorphosis, in which the first pair of parapodia are transformed into posterior tentacles of the head, occurs after the juveniles have begun to add their 6^th^ posterior segment [13–16].

Recently, we identified myoinhibitory peptide (MIP) as an inducer of rapid larval settlement behavior in *Platynereis* [11]. MIP is expressed in anterior chemosensory-neurosecretory neurons of the larva. Exogenous application of MIP inhibits the activity of the locomotor cilia, resulting in rapid sinking, and induces sustained contact with the substrate. *Platynereis* MIP belongs to an ancient neuropeptide family of Wamides, which are characterized by their amidated C-terminal tryptophan residue preceded by a small aliphatic residue [11, 17]. Wamides are widespread among eumetazoans, except deuterostomes, and recently emerged as conserved regulators of life cycle transitions [18]. For example, in some insects, MIP (also known as prothoracicostatic peptide (PTSP) or allatostatin-B (AST-B)) regulates ecdysone [19–21] and juvenile hormone levels [22], potentially influencing the timing of larval ecdysis and pupation. In cnidarians, including some corals and hydrozoans, GLWamide (also called metamorphosin) is known to induce larval settlement and metamorphosis [23–25].

How changes in feeding are regulated during marine life cycle transitions is less well understood. Many neuropeptides are known to have roles in regulating different aspects of feeding [26–28]. MIPs/Wamides are also pleiotropic [29–35] and regulate aspects of feeding and gut muscle activity in some insects and cnidarians. The first MIP described had a myoinhibitory function on adult locust hindgut [36]. In several insects, MIP is expressed in the adult stage and can suppress muscle contractions of the hindgut [36–40]. MIP is also expressed in the stomatogastric nervous system of the adult crab, *Cancer borealis*, where it decreases the frequency of pyloric rhythm [41, 42]. In addition, cnidarian GLWamide increases myoactivity in both hydra and sea anemone polyps, potentially influencing feeding [43, 44]. Although none of these studies directly quantified feeding in the whole organism, MIP is a strong candidate for the regulation of feeding during marine life-cycle transitions.

Here, we study the expression and function of MIP in *Platynereis* during late larval (3-6 days) and early juvenile development (6-30 days). We found MIP expression in sensory neurons of the gut of 6 days and older *Platynereis*. We used both peptide-soaking and morpholino-mediated knockdown approaches to establish a role for MIP in the regulation of postlarval feeding and gut peristalsis. MIP treatment also resulted in faster juvenile growth, probably as a consequence of increased food ingestion and gut movement. Our results establish MIP as a pleiotropic neuropeptide in *Platynereis* that links behavioral and physiological components of a life-cycle transition.

## Results

### *Platynereis MIP* continues to be expressed following larval settlement

Expression profiling of *MIP* precursor gene and its corresponding MIP peptide by RNA *in situ* hybridization and immunostaining respectively showed that MIP is expressed during both larval and postlarval development and continues to be expressed after cephalic metamophosis, in the early adult stage (Figure 1A-C, G-H; Additional File 1). At 6 days and older, *MIP* is expressed in both the median brain and the trunk nervous system, in paired cells and also in single cells closer to the larval midline. We also found MIP expression in the digestive system, both in the fore-and the hindgut. The *MIP*-expressing cells in the gut have sensory dendrites that project toward the lumen of the gut (Figure 1B-C; Additional Files 2, 3). In some of these dendrites we could even detect the *MIP* RNA *in situ* signal, allowing the unambiguous assignment of these acetylated tubulin-positive cellular projections to the MIP-expressing cells (Additional File 3).

**Figure 1.**
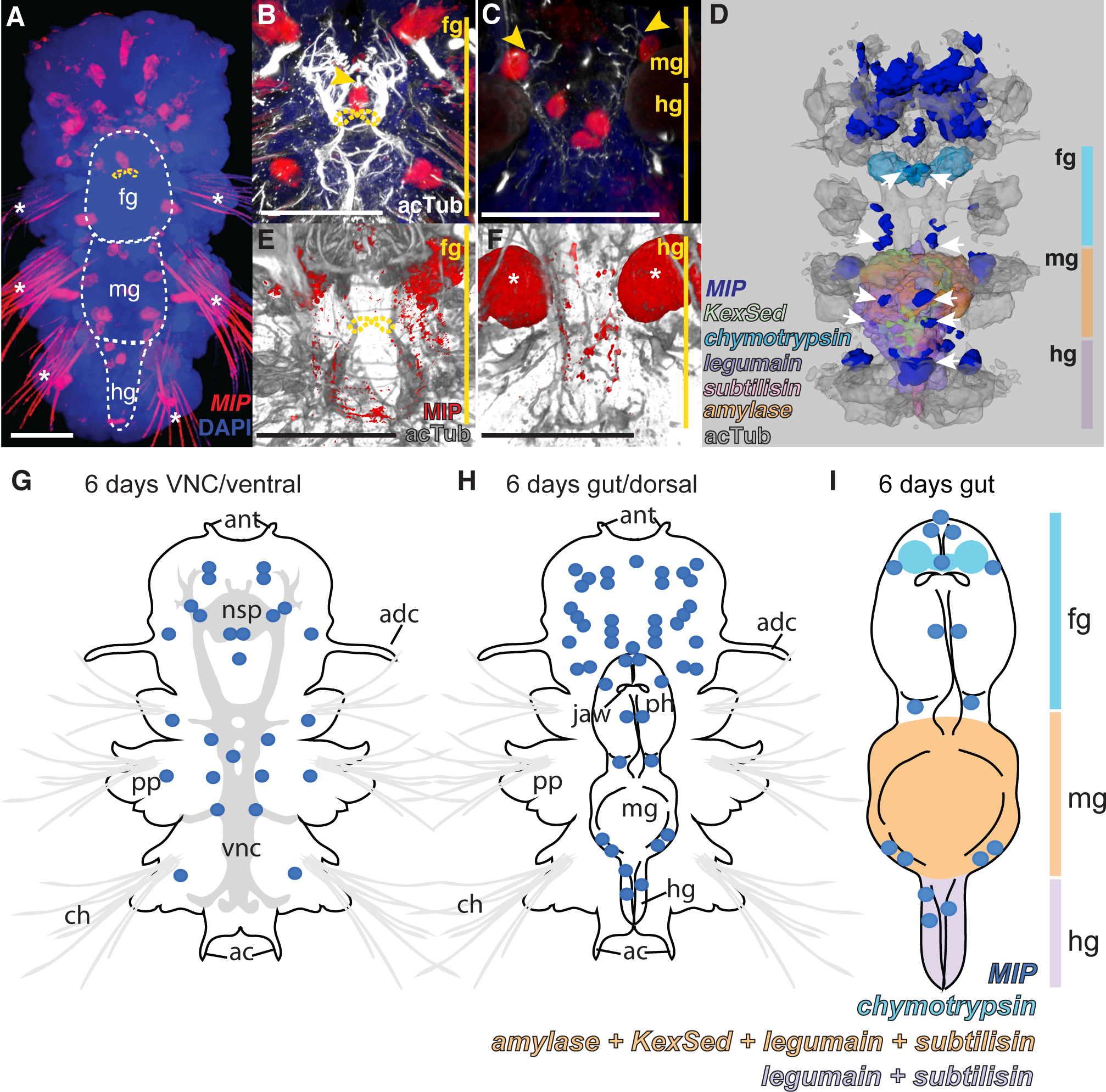
MIP is expressed in the digestive system of 6 day *Platynereis*. **(A)** Whole-mount RNA *in situ* hybridization (WMISH) for the *Platynereis MIP* precursor (red) counterstained with DAPI nuclear stain (blue). Ventral view of 6 day *Platynereis* with ventral nerve cord (VNC) region removed to reveal the underlying digestive system (outlined with white dashed line). **(B,C)** Close-up WMISH for the *Platynereis MIP* precursor (red) counterstained for acetylated tubulin (acTub) (white) and DAPI nuclear stain (blue). Yellow arrowheads indicate sensory dendrites of *MIP*-expressing cells. **(B)** Ventral view of the foregut. **(C)** Ventral view of the midgut and hindgut. Yellow dashed lines indicate jaws. White asterisks indicate background reflection from chaetae or spinning glands. **(D)** Dorsal view of surface representation of average *MIP* precursor expression domains registered to a 6 day acetlated tubulin reference template. Expression domains of digestive enzyme genes *chymotrypsin*, *kexin sedolisin* (*KexSed)*, *legumain*, *subtilisin* and *amylase* are included as digestive system markers. White arrows indicate areas of *MIP* expression associated with the digestive system. **(E,F)** Immunostaining with *Platynereis* MIP antibody (red) counterstained with acetlated tubulin (grey) at 6 days, with VNC removed to show digestive system. **(E)** Ventral view of foregut. **(F)** Ventral view of hindgut. **(G)** Schematic of *MIP* precursor expression on ventral side of 6 days *Platynereis*. **(H)** Schematic of *MIP* precursor expression on dorsal side of 6 days *Platynereis*. **(I)** Schematic overview of expression of *MIP* and digestive enzymes in the gut of 6 day *Platynereis*. Scale bars: 50 µM. Abbreviations: fg, foregut; mg, midgut; hg, hindgut; ant, antenna; nsp, neurosecretory plexus; adc, anterior dorsal cirrus; pp, parapodia; vnc, ventral nerve cord; ch, chaetae; ac, anal cirrus; ph, pharynx.

Immunostaining with the *Platynereis* MIP antibody showed that in addition to the neurosecretory plexus of the brain, MIP peptide is transported throughout the ventral nerve cord. At 6 days, MIP-expressing neurons innervate both the foregut and hindgut (Figure 1E,F). As larvae progress from 3 days to 6 days, during which time the digestive system develops, MIP expression first emerges in the hindgut, followed by the expression in the foregut (Additional File 4A-L). By one month, MIP-expressing cells densely innervate the entire length of the gut forming a nerve-net. Using an antibody against the conserved C-amidated dipeptide VWamide [11], we found similar immunolabeling in the brain, ventral nerve cord and gut of larvae of *Capitella teleta*, a distantly related annelid species [45] (Additional File 4M-P). These results suggest a potential role for MIP signaling in feeding and digestion during larval and early juvenile stages of the life cycle.

### Characterization of normal gut development and the initiation of feeding in postlarvae

At 6 days the *Platynereis* postlarvae have a through gut with clearly recognizable fore-, mid-and hindgut regions (Figure 1). The foregut contains the muscular and extendable pharynx with the jaws and salivary glands (Additional file 2). The foregut-midgut boundary is marked by the presence of a sphincter muscle. This muscle showed regular contractions in larvae expressing a genetically encoded calcium indicator GCaMP6 (Additional File 5). The broad midgut does not show regionalization and is followed by a short and narrow hindgut.

To gain insight into the morphology and maturation of the *Platynereis* digestive system, we carried out wholemount RNA *in situ* hybridization on 6 day, 14 day and 1 month samples with digestive enzyme genes: peptidases *kexin sedolisin*, *subtilisin*, *chymotrypsin* and *legumain*, and the polysaccharide-digesting enzyme *alpha*-*amylase* (Figure 1D,I; Additional Files 6-8). *Alpha-amylase* and *subtilisin* expression was restricted to the midgut at 6 days, but expanded to mid-and hindgut at 14 days and 1 month. *Legumain* was constantly expressed in both mid-and hindgut from 6 days to 1 month, while *kexin-sedolisin* expression was restricted to the midgut from 6 days to 1 month. *Chymotrypsin alpha* was the only digestive enzyme gene tested with expression in the foregut, including the salivary glands, at 6 and 14 days. At 1 month, *chymotrypsin alpha* remained strongly expressed in the foregut, but expression also extended to the mid-and hindgut. Registration of these digestive gene expression patterns [46] at 6 days to a common acetylated tubulin reference scaffold along with the average 6 day *MIP* expression highlighted the close association of *MIP*-expressing cells with the digestive system at this stage (Figure 1D; Additional File 8).

We also looked at the change in expression of these digestive enzyme genes and *MIP* across the *Platynereis* life cycle in stage-specific RNA-seq datasets [47]. With the exception of *legumain*, the expression of all digestive enzymes was undetectable in non-feeding larval stages but sharply increased between 4 and 10 days (Additional File 9). In accordance with their digestive function, these enzymes were strongly down-regulated in the adult non-feeding epitokes. *MIP* expression also increased sharply between 4 and 10 days, although it continues to be expressed in the non-feeding epitokes, suggesting further functional roles beyond feeding in *Platynereis*. Following settlement from the water column, *Platynereis* larvae have been reported to begin feeding between 5 – 8 days, with considerable variation between individuals [13, 16]. Due to this variability, we decided to document feeding initiation in our own laboratory culture. We added *Tetraselmis marina* microalgae to the larval cultures and documented feeding based on chlorophyll fluorescence in the gut. Most larvae initiated feeding between 6 and 7 days; by 8 days, nearly all larvae had started feeding (Figure 2A, Figure 3C).

**Figure 2.**
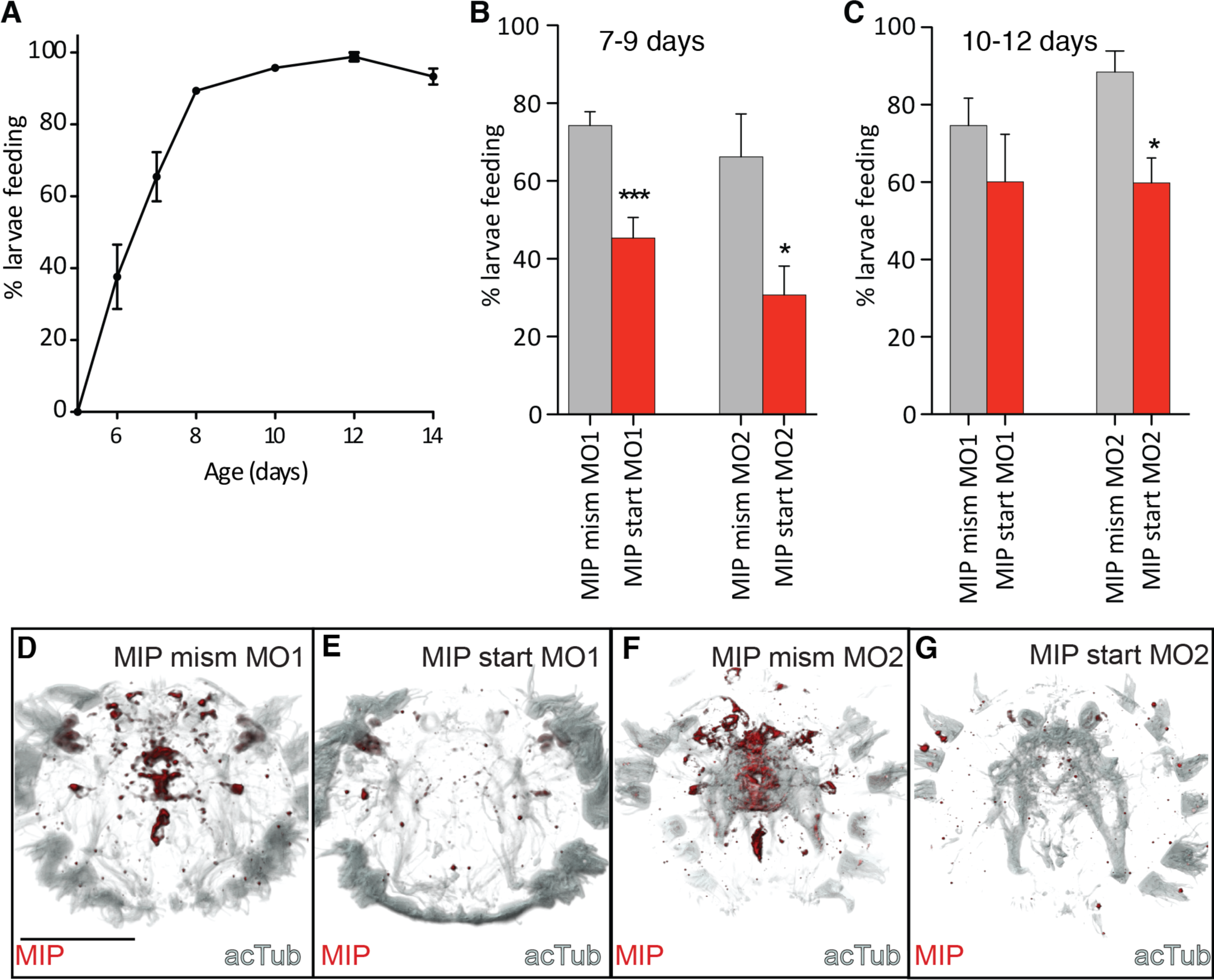
Feeding in control and MIP knockdown larvae. **(A)** Timecourse of initiation of feeding during *Platynereis* larval development. % larvae feeding on *Tetraselmis marina* algae between 5 – 14 days. Data are shown as mean +/− s.e.m. **(B)** % larvae feeding at 7-9 days following injection of MIP mismatch or start morpholinos **(C)** % larvae feeding at 10-12 days following injection of MIP mismatch and start morpholinos. Data are shown as mean +/− s.e.m. p-value cut-offs based on unpaired *t*-test: *** <0.001; ** < 0.01; * <0.05. **(D–G)** Anterior view of 6 day *Platynereis* injected with MIP mismatch or start morpholinos and immunostained with *Platynereis* MIP antibody (red) counterstained with acetylated tubulin (grey). **(D)** Larva injected with mismatch morpholino 1 **(E)** Larva injected with MIP start morpholino 1 **(F)** Larva injected with mismatch morpholino 2 **(G)** Larva injected with MIP start morpholino 2. Identical confocal microscopy and image processing parameters were applied to all images. Scale bar: 100 µm. Abbreviations: mism, mismatch; MO, morpholino; acTub, acetylated tubulin.

**Figure 3.**
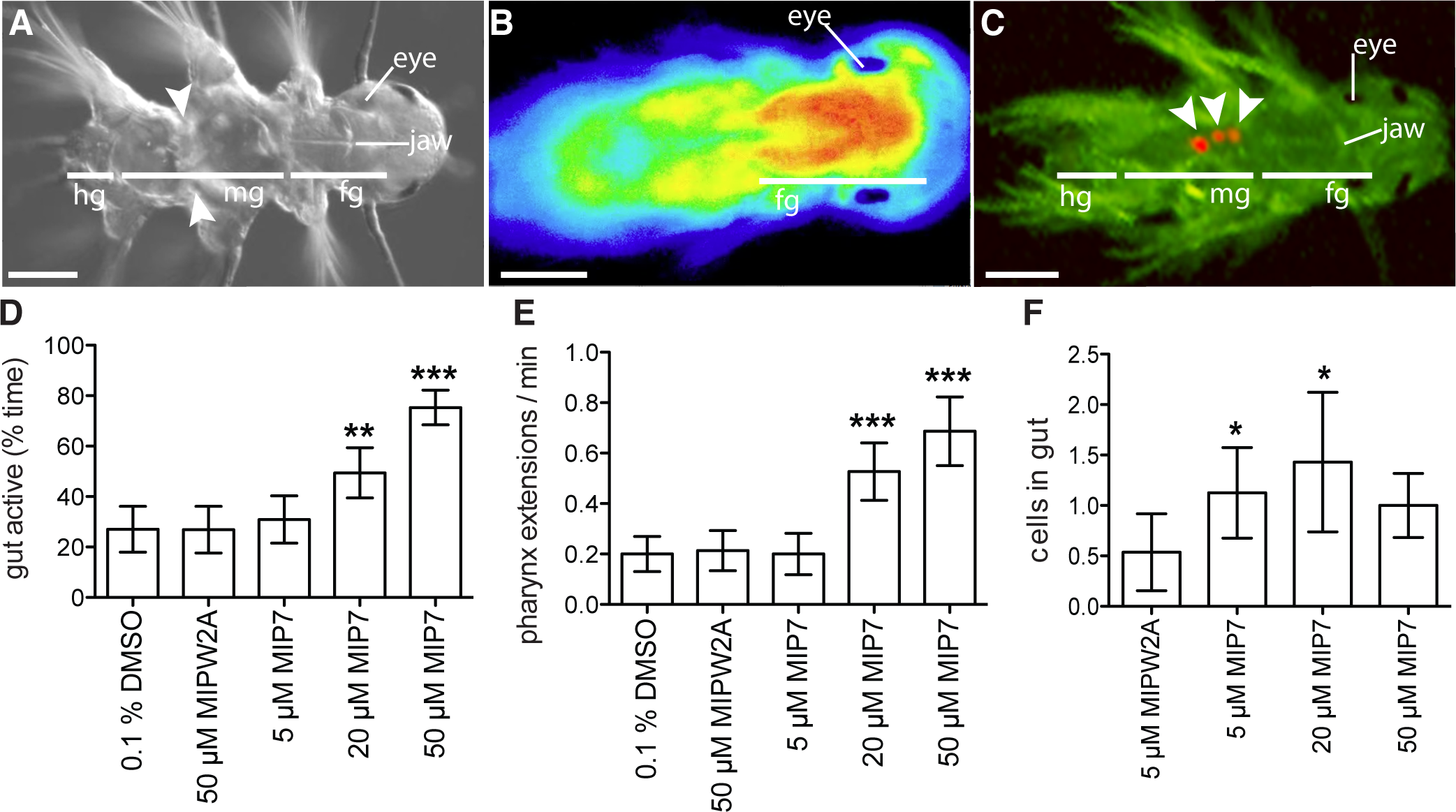
Synthetic MIP treatment increases gut peristalsis, pharynx extension and ingestion in *Platynereis*. **(A)** Differential interference contrast micrograph of 6.5 day *Platynereis*. White arrowheads indicate muscular contraction in the gut. **(B)** Calcium-imaging with GCaMP6 in 6.5 day *Platynereis* highlights muscular pharynx extension in the foregut. **(C)** Fluorescent micrograph of 7 day *Platynereis* with AF488 filter. White arrowheads indicate autofluorescent *Tetraselmis* cells in the gut. All images are dorsal views, with head to the right. **(D)** Gut activity as percentage of time in MIP-treated versus control 6.5 day *Platynereis*. **(E)** Number of pharynx extensions per minute in MIP-treated versus control 6.5 day *Platynereis*. **(F)** Number of *Tetraselmis marina* algae cells eaten per larvae in 30 min in MIP-treated versus control 7 day *Platynereis*. Data are shown as mean +/− 95% confidence interval, n = 60 larvae. p-value cut-offs based on unpaired *t*-test: *** <0.001; ** < 0.01; * <0.05. MIPW2A is a control non-functional MIP peptide in which the two conserved tryptophan sites are substituted with alanines. Scale bars: 50 µm. Abbreviations: hg, hindgut; mg, midgut; fg, foregut.

### Knockdown of MIP delays the initiation of feeding in *Platynereis* larvae

To explore the function of MIP in the *Platynereis* digestive system, we employed morpholino microinjection to knockdown MIP expression. We used two different translation blocking morpholinos and two mismatch control morpholinos. To test the effectiveness of MIP-knockdown, we immunostained knockdown and control larvae with an antibody against *Platynereis* MIP. We observed a strong reduction in MIP immunostaining in *Platynereis* MIP-knockdown larvae, but not in controls, up to at least 6 days (Figure 2D-G). These experiments confirmed that the MIP morpholinos were capable of strongly reducing MIP expression.

Next, we documented feeding in MIP-knockdown and control larvae. Similar to untreated larvae, most larvae injected with a control morpholino had initiated feeding between 7-9 days, whereas a significantly lower number of MIP-knockdown larvae had food in the gut at 7-9 days. This effect was still observed between 10-12 days (Figure 2 B-C).

To rule out that the reduced feeding in MIP-knockdown larvae is due to a developmental delay, we compared the morphology of the nervous system of MIP-knockdown and control larvae. There were no detectable differences in the nervous system of control larvae and MIP-knockdown larvae based on acetylated tubulin immunostainings at 6 days (Figure 2D-G; Additional File 10 A, B). We also treated uninjected larvae between 24 h and 5 days with synthetic MIP peptide to see whether MIP-treated larvae initiate feeding sooner, indicating a potential developmental acceleration. MIP treatment did not significantly alter the timing of feeding initiation, even when food was available earlier than 5 days (Additional file 11).

These knockdown experiments establish a critical physiological role for MIP in the initiation of feeding in *Platynereis* postlarvae.

### MIP treatment has a myostimulatory effect on the digestive system of *Platynereis* postlarvae

In order to understand how the morpholino knockdown of MIP resulted in reduced larval feeding, we examined the effect of synthetic MIP treatment on postlarvae with a focus on the digestive system. Treatment of 6.5 day postlarvae with synthetic MIP caused a significant increase in gut peristalsis (Figure 3A, D; Additional File 12). MIP-treated postlarvae also displayed increased rates of pharynx extension (Figure 3E; Additional File 13).

To determine whether these effects on gut and pharynx movement resulted in increased ingestion of algal cells in MIP-treated postlarvae, we then scored the number of algal cells consumed by MIP-treated versus control 7 day postlarvae. Treatment with 5 µM and 20 µM MIP significantly increased postlarval algal cell consumption (Figure 3C,F; Additional File 14). Additionally, MIP-treated postlarvae have decreased crawling speed, indicating a switch in the nervous system from a locomotory to a feeding program (Additional File 15). These results show that MIP directly up-regulates feeding activity and gut peristalsis.

### Long-term MIP treatment enhances growth in *Platynereis* postlarvae

At approximately two weeks of age, feeding *Platynereis* begin to add new posterior segments [13]. After the development of the 5^th^ segment juveniles undergo cephalic metamorphosis, a morphogenetic process in which the first chaetigerous segment loses its chaetae, develops a pair of tentacular cirri and fuses with the head (Figure 4A-D). The timing of cephalic metamorphosis and the addition of new segments varies between individuals. Even juveniles cultured individually showed variation in the timing of posterior segment addition, with segment number varying between 4 and 8 segments at 34 days (Additional File 16). On a diet of *Tetraselmis*, the shortest interval for an individually-raised juvenile to develop an additional posterior segment was 4 days. The addition of new segments is strictly dependent on feeding. Unfed larvae never develop beyond the 3-segmented stage (Additional File 16). Growth in other nereid species depends on culture density [48–51]. We documented the growth of *Platynereis* juveniles cultured at different densities with excess food and determined the maximal density that still allowed optimal growth (3 larvae/ml)

**Figure 4.**
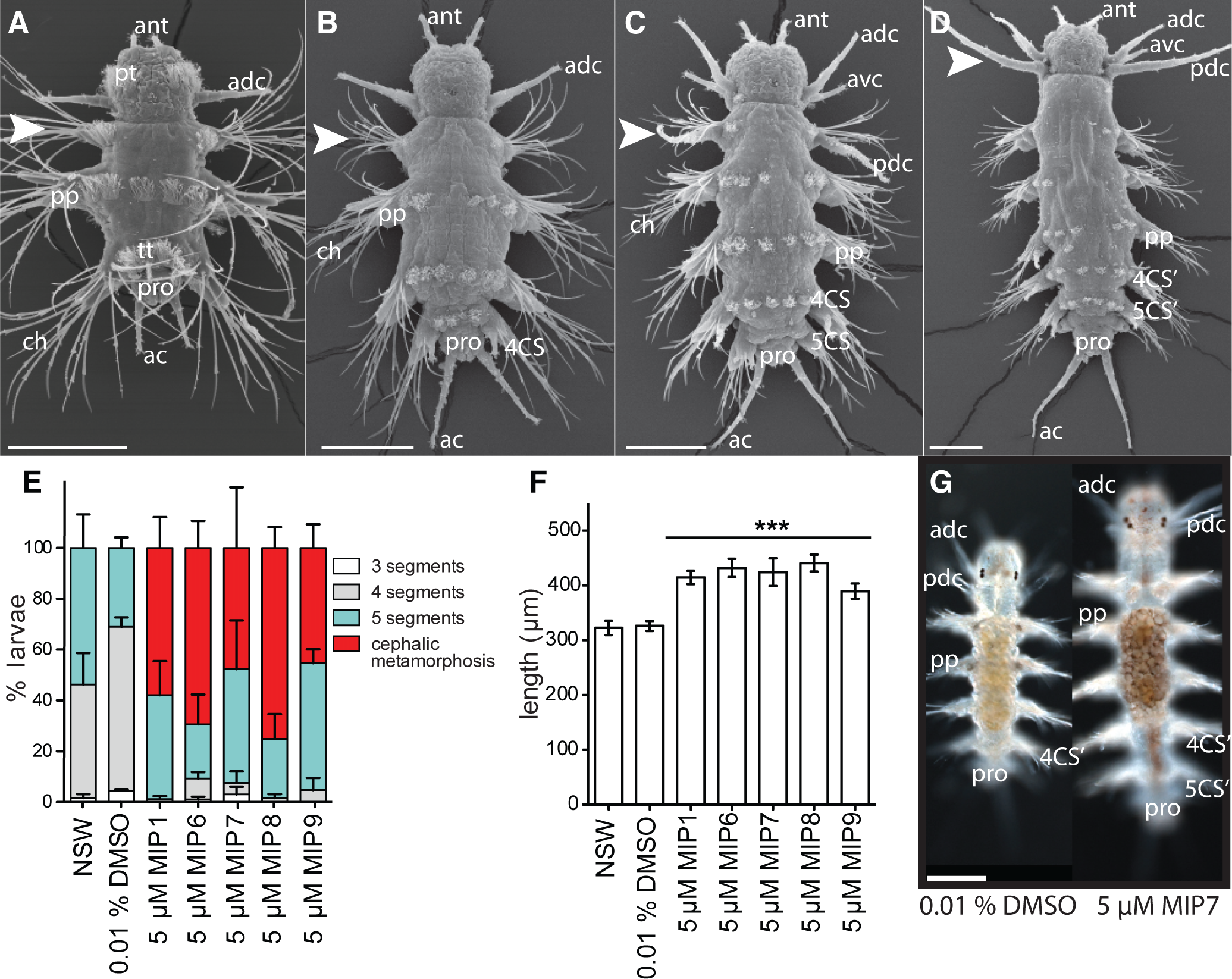
Long-term treatment of *Platynereis* with synthetic MIP enhances growth and decreases time to cephalic metamorphosis. (A-D) SEM images depicting posterior segment addition followed by cephalic metamorphosis in *Platynereis dumerilii*, dorsal views. White arrowhead indicates parapodia of the 1^st^ chaetigerous segment, which are transformed into the posterior tentacular cirri of the head during cephalic metamorphosis. (A) 6 day 3-segmented nectochaete larva. (B) 4-segmented errant juvenile. (C) 5-segmented errant juvenile. Cephalic metamorphosis has begun. (D) Cephalic metamorphosis is complete. (E) Percentage of 25 day *Platynereis* larvae/juveniles with 3, 4, 5 segments, or complete cephalic metamorphosis following exposure to 5 µM synthetic MIP peptides, from 4 days onwards. Data are shown as mean +/− s.e.m. NSW, natural seawater control. (F) Total length of 25 day *Platynereis* larvae/juveniles exposed to 5 µM synthetic MIP peptides, from 4 days onwards. Data are shown as mean +/− 95% confidence interval. p-value cut-offs based on unpaired *t*-test: *** <0.001; ** < 0.01; * <0.05. **(G)** Differential interference contrast light micrographs of example control and MIP7-treated individuals at 1 month. Scale bars: 100 µM. Abbreviations: ac, anal cirrus; adc, anterior dorsal cirrus; ant, antenna; avc, anterior ventral cirrus; ch, chaetae; pdc, posterior dorsal cirrus; pp, parapodia; pro, proctodeum; pt, prototroch; tt, telotroch; 4CS, 4^th^ chaetigerous segment; 5CS, 5^th^ chaetigerous segment; 4CS’, 4^th^ chaetigerous segment after cephalic metamorphosis; 5CS’, 5^th^ chaetigerous segment after cephalic metamorphosis.

(Additional File 16). Under these conditions, juvenile *Platynereis* begin to develop the 5^th^ segment at 16 days, and start to undergo cephalic metamorphosis at 24 days. Morpholino knockdown methods are not applicable to such late stage animals, therefore we tested the effects of MIP treatment on errant juvenile growth. We found that the time to the addition of new posterior segments, and to cephalic metamorphosis was reduced by sustained exposure to different versions of MIP (Figure 4E). At 25 days, some MIP-treated individuals had completed cephalic metamorphosis, while control individuals were yet to undergo cephalic metamorphosis. The effect of MIP treatment on growth was only seen in the presence of food. In the absence of food, MIP treatment could not induce the addition of any new posterior segments, and postlarvae remained at the 3-segmented stage (Additional File 16). Additionally, MIP-treated larvae, both fed and unfed, exhibited altered pigmentation of the gut and the body (Figure 4G,H, Additional File 16).

## Discussion

Our results established the MIP neuropeptide as a regulator of postlarval feeding and gut activity in *Platynereis*. At 6 days, MIP is expressed in both the pharynx and the hindgut in neurons with a sensory morphology with dendrites projecting to the lumen. This suggests that MIP cells receive sensory signals from inside the mouth and the gut and respond by releasing MIP in a neurosecretory manner in the vicinity of the pharynx and hindgut muscles. Contrary to its name, the local release of MIP could have a myostimulatory role and increase the rate of pharynx extensions and peristaltic movements. Alternatively, MIP may regulate other neurons in the gut, for example, in an as yet unidentified central pattern generator circuit responsible for regular gut contractions, as seen in crustaceans [52]. Such a direct sensory-neuromodulatory mechanism could contribute to the initiation of the feeding program in postlarvae following settlement upon the first encounter with food.

Our results show that the increase in pharynx extensions in *Platynereis* postlarvae has a direct effect on the amount of food ingested. Increased gut peristalsis could promote the passage of food within the gut or the mixing of food with digestive enzymes, speeding up digestion. The sustained expression of MIP in the gut and the long-term effects of MIP on juvenile growth indicate that MIP also has an important physiological role later in the life cycle. The enhancement of juvenile growth we interpret as a consequence of increased feeding. Given its effect on both settlement and growth, MIP treatment may be a useful means of enhancing both larval settlement and juvenile growth in polychaete aquaculture [53].

However, the morpholino knock-down experiments show that without MIP postlarvae start feeding later.

Interestingly, MIP has a myostimulatory role in cnidarians and *Platynereis*, but a myoinhibitory role in arthropods [36, 41, 44]. This could mean that either MIP was independently recruited to regulate gut activity in different lineages, or that the sign of the regulation switched during evolution. In the latter case, MIP would represent a conserved bilaterian gut peptide influencing feeding. Further comparative morphological and molecular studies of MIP cells and signaling pathways in a broader range of taxa will be needed to resolve this.

MIP regulates both settlement behavior and feeding, two aspects of the pelagic-to-benthic transition of the non-feeding *Platynereis* larvae. What could be the reason for the redeployment of the same peptidergic signal at different times during development and in different contexts? One possibility is that the anterior MIP-expressing sensory-neurosceretory cells of the larva and the MIP cells in the gut of the postlarva sense the same chemical cues released by potential food sources. Some marine larvae are induced to settle by their future juvenile food source [2]. Testing this hypothesis will require the identification of settlement cues and their corresponding receptors in *Platynereis*.

In *Platynereis*, juvenile feeding is an essential requirement for the completion of cephalic metamorphosis. In other polychaete species, where feeding often begins in the pelagic larval stage before settlement, feeding is also an essential component for settlement and metamorphosis. Starved larvae of *Capitella sp*., *Polydora ligia*, *Hydroides elegans* and *Phragmatopoma lapidosa* all lose or have decreased ability to complete settlement and metamorphosis [54–57]. Exploration of the roles of MIP in polychaete species with feeding larvae would increase our understanding of the links between MIP signaling, larval settlement and feeding.

## Conclusions

We have described a role for MIP in *Platynereis* postlarval feeding and established methods for studying the neuroendocrine regulation of feeding, providing the basis for future studies in this area. The amenability of *Platynereis* larvae to peptide treatments by soaking, the transparent body and an ancestral neuropeptide complement make *Platynereis* an ideal model with which to study the neuroendocrine regulation of feeding.

## Methods

### Platynereis culture

*Platynereis* larvae were obtained from an in-house culture as previously described [15]. After fertilization of eggs, developing embryos and larvae were kept in an incubator at a constant temperature of 18°C with a regular light-dark cycle.

### RNA *In situ* hybridization

Different developmental stages of *Platynereis* were collected for fixation for use in wholemount RNA *in situ* hybridization and immunostaining techniques. Individuals 6 days and older were relaxed using 1 M MgCl_2_ [58] prior to fixation. Postlarvae and juveniles that had begun feeding were starved for a few days prior to fixation to avoid the presence of autofluorescent algae cells in the gut, which interfere with fluorescent signals from immunostaining. All animals were fixed in 4% paraformaldehyde in 0.1 M MOPS (pH 7.5), 2 mM MgSO_4_, 1 mM EGTA, 0.5 M NaCl for 1 h at room temperature. Fixed larvae were dehydrated through a MeOH series and stored in 100% MeOH at −20°C.

DIG-labelled antisense RNA probes for the *Platynereis-MIP* precursor, *alpha-amylase*, *subtilisin, legumain, kexin-sedolisin* and *chymotrypsin alpha* were synthesized from purified PCR products of clones sourced from a *Platynereis* cDNA library [47]. Digestive enzyme genes were identified by the presence of a signal peptide assigned using the SignalP 4.1 server (http://www.cbs.dtu.dk/services/SignalP/) and conserved domains assigned by a BLASTP against the Pfam database with e-value cutoff 1e-06 (http://pfam.xfam.org/search). RNA *in situ* hybridization using nitroblue tetrazolium (NBT)/5-bromo-4-chloro-3-indolyl phosphate (BCIP) staining combined with mouse anti-acetylated-tubulin staining to highlight cilia and nervous system, followed by imaging with a Zeiss LSM 780 NLO confocal system and Zeiss ZEN2011 Grey software on an AxioObserver inverted microscope, was performed as previously described [46], with the following modification: fluorescence (instead of reflection) from the RNA *in situ* hybridization signal was detected using excitation at 633 nm in combination with a Long Pass 757 filter. Animals were imaged with a 40X oil objective.

### Image Registration of RNA *in situ* hybridization patterns

We projected average *MIP* expression pattern of four 6 day individuals onto a common 6 day whole-body nuclear reference template generated from DAPI signal of forty 6 day individuals as described previously for 72 h larvae [46]. Acetylated tubulin and expression patterns of digestive system marker genes of select individuals were also projected onto the reference template. Snapshots and video of the projected genes were generated in Blender 2.7.1 (http://www.blender.org/).

### Immunostaining

Rabbit antisera against *Platynereis* MIP7 were affinity-purified using the synthetic peptide coupled to a Sulfolink resin (Thermo Scientific) via an N-terminal Cys (CAWNKNSMRVWamide) as previously described [59]. Immunohistochemistry with 1 µg/ml rabbit MIP7 and 0.5 µg/ml mouse anti-acetylated tubulin (Sigma) primary antibodies, and 1 µg/ml anti-rabbit Alexa Fluor^®^ 647 (Invitrogen) and 0.5 µg/ml anti-mouse FITC (Jackson Immuno Research) secondary antibodies, was performed as previously described [59]. Images were processed with Imaris 6.4 (Bitplane Inc., Saint Paul, USA) software. Raw image stacks are available at the Dryad data repository.

### Calcium Imaging

Fertilized eggs were injected with 500 ng/µl capped and polyA-tailed GCaMP6 [60] RNA generated from a vector (pUC57-T7-RPP2-GCaMP6) containing the GCaMP6 ORF fused to a 169 base pair 5’ UTR from the *Platynereis* 60S acidic ribosomal protein P2, as in [61]. Injection protocol is described in more detail in the ‘Morpholino Knockdown of *Platynereis* MIP’ section. Larvae were imaged with a 488 nm laser and transmission imaging with DIC optics on a Zeiss LSM 780 NLO confocal system on an AxioObserver inverted microscope (Additional File 5), or using a Zeiss AxioZoom.V16 microscope with Hamamatsu Orca-Flash 4.0 digital camera (Additional File 13).

### RNA-Seq

RNA-Seq analysis of digestive enzyme and *MIP* precursor gene expression was performed on an existing dataset of 13 different stages spanning the *Platynereis* life cycle, from egg to mature adults. Methods used were described in [47].

### Documentation of normal feeding in *Platynereis*

To document variation in the commencement of feeding in *Platynereis* larvae from our laboratory culture, larvae were kept in Nunclon 6-well tissue culture dishes, with 10 ml sterile filtered seawater (FSW) per well. Each well contained 30 larvae. Larvae from 6 different batches, with different parents, were used in our analysis. Larvae were fed 5 µl *Tetraselmis marina* algae culture at 5 days. Larvae were then tested for feeding by checking for the presence of fluoresent *Tetraselmis* algae in the gut using a Zeiss Axioimager Z1 microscope with an AF488 fluorescent filter and a 20X objective. Larvae were checked for signs of feeding at 5.5, 6, 7, 8, 10, 12 and 14 days. After ingestion, algae cells can remain in the gut for up to 48 h before digestion causes a loss of fluorescence. Although larvae with a full gut can also be identified with normal light microscopy due to the transparent body wall, fluorescent microscopy enables the detection of even a single alga cell in the gut, due to the strong chlorophyll fluorescence of the *Tetraselmis* cells.

### Morpholino knockdown of *Platynereis* MIP

Two translation blocking morpholinos (MOs) and two corresponding 5 base pair mismatch control morpholinos were designed to target the *Platynereis-MIP-precursor* (GeneTools, LLC):

Pdu-MIP-start MO1 TGATAGTGACGCGATCCATTGGACT

Pdu-MIP-mism MO1 TGTTAGTGACCCGTTCGAATGGACT

Pdu-MIP-start MO2 CTAGTTCCTTCTCTCCCTCTTATCT

Pdu-MIP-mism MO2 CTACTTGCTTGTCTCCGTGTTATCT

Nucleotides complementary to the start codon (ATG) are underlined; nucleotides altered in mismatch control morpholinos are highlighted in grey.

MOs were diluted in water with 12 µg/µl fluorescein dextran (*Mr* 10,000, Invitrogen) as a fluorescent tracer. 0.6 mM MOs were injected with an injection pressure of 600 hPa for 0.1 s and a compensation pressure of 35 hPa using Eppendorf Femtotip II needles with a Femtojet microinjector (Eppendorf) on a Zeiss Axiovert 40 CL inverted microscope equipped with a Luigs and Neumann micromanipulator. The temperature of developing zygotes was maintained at 16°C throughout injection using a Luigs and Neumann Badcontroller V cooling system and a Roth Cyclo 2 water pump.

For microinjection, fertilized *Platynereis* eggs developing at 16°C were rinsed 1 h after fertilization with sterile 0.2 µm filtered seawater (FSW) in a 100 µM sieve to remove the egg jelly, followed by a treatment with 70 µg/ml proteinase K for 1 min to soften the vitellin envelope. Following injection, embryos were raised in Nunclon 6-well plates in 10 ml FSW and their development was monitored daily.

Larvae were fed 5 µl *Tetraselmis marina* algal culture at 6 days. Feeding in 7 – 14 day injected larvae was assessed by checking for the presence of fluoresent *Tetraselmis marina* algae in the gut using a Zeiss Axioimager Z1 microscope with an AF488 fluorescent filter and a 20X objective. Larvae were checked for signs of feeding as described above every 24 h from 7 days on. We scored a minimum of 62 larvae (maximum 109 larvae) from a minimum of 3 separate microinjection sessions (with 3 different batches of larvae) for each translation-blocking and control morpholino. Photomicrographs of morpholino-injected larvae were also taken and larval body length was measured from these pictures using Image J 64 software. Some morpholino-injected larvae were also fixed at 6 days for immunostaining with the anti-MIP antibody (as described above) in order to assess morpholino specificity and effectiveness.

### Effect of synthetic MIP on *Platynereis* feeding behaviour

Peptide functions can be investigated in *Platynereis* larvae by bath application of synthetic neuropeptides [10, 11]. To test whether synthetic MIP treatment increased developmental speed, leading to early initiation of feeding in *Platynereis* larvae, experiments were performed in Nunclon 6-well plates, with 10 ml FSW per well. Each control and peptide treatment was replicated across three wells, with 30 larvae per well. Larvae were treated with 5 µM MIP7 or controls at 24 h, 60 h, 4 days or 5 days, then fed at 4 or 5 days (depending on the age at which MIP treatment occurred) with 5 µl *Tetraselmis marina* algal culture. Larvae were fed at an earlier age due to the possibility of MIP treatment causing an earlier initiation of feeding. Larvae were checked for feeding by monitoring algal cell fluorescence in the gut as described above. Larvae were monitored from 5 or 5.5 days (depending on age at which larvae were first fed) until 7 or 8 days. A control non-functional MIP peptide (MIPW2A, AANKNSMRVAamide), in which the two conserved tryptophan sites were replaced with alanines (this prevents MIP from binding to its receptor, see [11]) was also tested. A further control of larvae treated with DMSO alone was also included, as MIP peptides require DMSO to be dissolved in solution.

To test the effects of synthetic MIP peptide treatment on the digestive system of *Platynereis* larvae, we recorded videos of groups of 60 larvae in a square glass cuvette 1.5 cm x 1.5 cm x 0.3 cm in 500 µl of FSW using a Zeiss AxioZoom.V16 microscope with Hamamatsu Orca-Flash 4.0 digital camera. For each treatment and control, 3 biological replicates (larval batches with different parentage, fertilized on different days) were carried out. We tested three concentrations of synthetic MIP: 5, 20 and 50 µM, plus 50 µM control non-functional MIP peptide MIPW2A and a 0.1 % DMSO control. A 2.5 min video at 10 frames per second was recorded 10 min after peptide or DMSO addition. Videos were analyzed manually in Fiji (Image J 1.48s, Wayne Rasband, http://imagej.nih.gov/ij). For each video, 20 larvae that remained within the frame of the video for the entire 2.5 min were scored for gut peristalsis and pharynx extension activity. Distance traveled and speed of the larvae was also measured using the MTrack2 plugin [62]. Significant differences in gut peristalsis, pharynx extension activity and locomotion in MIP-treated versus control larvae were tested in an unpaired *t* test.

To test the effects of synthetic MIP treatment on short-term ingestion of algal cells in *Platynereis* larvae, experiments were performed in Nunclon 24-well plates, with 2 ml FSW per well. Each control and peptide treatment was replicated across three wells, with 20 larvae per well. 7 day postlarvae were treated with 5 µM MIPW2A control peptide, 5 µM MIP, 20 µM MIP or 50 µM MIP for 10 min. Following this, 20 µl *Tetraselmis marina* algal culture was added to each well and larvae were left to feed for 30 min. All larvae were then immediately fixed in 0.5 mL 4% paraformaldehyde in 1X PBS with 0.01% Tween (PTw) for 1 hour. Following 4 washes in 1 ml PTw, larvae were mounted on glass slides and the number of algal cells in the digestive system of each larvae was counted using a Zeiss Axioimager Z1 microscope with an AF488 fluorescent filter and a 20X objective. Significant differences in MIP-treated versus control larvae were tested in an unpaired *t* test.

### Scanning electron microscopy (SEM)

*Platynereis* larvae and juveniles of different developmental stages were fixed with 3% glutaraldehyde in 0.1 M phosphate buffer pH 7.2, rinsed in phosphate buffer, further fixed with 1 % osmium tetroxide in water and dehydrated in an ascending EtOH series over several days. Critical point drying with carbon dioxide was performed in a Polaron E 3000. The samples were coated with gold-palladium in a Balzers MED 010. Images were taken on a Hitachi S-800 Scanning electron microscope.

### Calculation of optimal culture density

The assessment of growth in larvae cultured individually was performed in a Nunclon 24-well tissue culture dish with 1 larva per well in 2 ml FSW. Larvae were fed from 6 days with 3 µl *Tetraselmis marina* algae culture. Larvae were scored under a dissection microscope for number of segments and cephalic metamorphosis every 48 h from 14 days to 34 days.

Documentation of growth in larvae cultured at different densities was carried out in Nunclon 6-well plates with 10 mL FSW/well and 30, 50 or 100 larvae per well. Three replicate wells were included for each culture density. Larvae were fed with surplus *Tetraselmis marina* algae culture throughout the experiment. Larvae were scored for segment number and cephalic metamorphosis every 4 days from 16 to 32 days.

## Long-term treatment of *Platynereis* with synthetic MIP

To test the effect of synthetic MIP treatment on growth in *Platynereis* larvae, we again carried out experiments in Nunclon 6-well plates as described above, with 30 larvae per well and 3 replicate wells per treatment and control. 5 µM synthetic peptides were added at 4 days. Different versions of mature MIP peptide tested were: MIP1 – AWNKNNIAWamide, MIP6 – AWGDNNMRVWamide, MIP7 – AWNKNSMRVWamide, MIP8 – AWKGQSARVWamide, and MIP9 – GWNGNSMRVWamide. Larvae were also fed at 4 days with 5 µl *Tetraselmis sp.* algal culture. At 25 days (21 days after peptide addition), errant juveniles were scored for number of segments and cephalic metamorphosis (as above). Juveniles were also photodocumented using a Zeiss Axioimager Z1 microscope with differential interference contrast (DIC) and size of control and treated larvae (end of head to end of pygidium, excluding cirri) was measured in Fiji (Image J 1.48s, Wayne Rasband, http://imagej.nih.gov/ij).

We also tested the effects of synthetic MIP treatment on growth of unfed *Platynereis* larvae. Experiments were carried out as above, 5 µM MIP7 was added at 4 days. 5 µM MIPW2A and 0.1% DMSO were included as negative controls. At 24 days, errant juveniles were scored for number of segments and cephalic metamorphosis. Most unfed juveniles died between 24 and 28 days.

## Competing interests

A patent application for the potential use of MIP/allatostatin B in lophotrochozoan larval culture has been submitted.

## Authors' contributions

EAW and GJ conceived and designed the study. EAW performed expression analysis, morpholino knockdown and behavioral/growth assays. MC performed expression analysis. EAW, MC and GJ analyzed the data. EAW drafted the manuscript. EAW, GJ and MC revised the manuscript. All authors read and approved the final manuscript.

## Acknowledgements

The authors thank Dorothee Hildebrandt for her assistance in caring for our *Platynereis* laboratory culture, Jürgen Berger for contributions to SEM, and Günter Purshke for sharing his knowledge of polychaete digestive systems. Aurora Panzera, Christian Liebig and Csaba Verasztó provided assistance with calcium imaging and microscopy. Reza Shahidi and Albina Asadulina assisted with image and video construction in Blender. Heiko Müller provided *in situ* probes for digestive enzyme genes. Nadine Randel provided fixed 6 day *Platynereis* larvae for SEM. The research leading to these results received funding from the European Research Council under the European Union’s Seventh Framework Programme (FP7/2007-2013)/European Research Council Grant Agreement 260821.

## Additional files

**Additional file 1.**
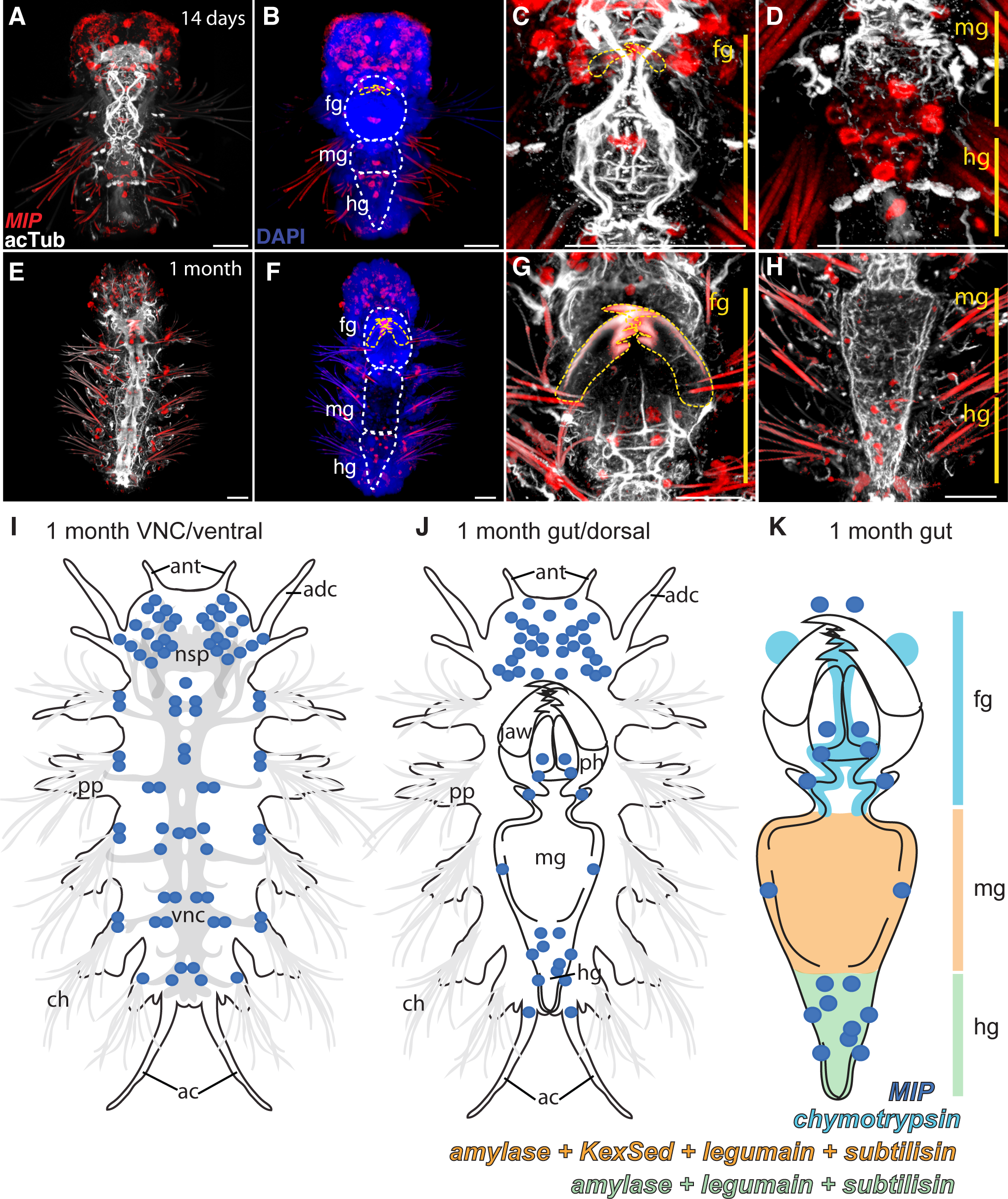
MIP expression in 14 day and 1 month *Platynereis*.pdf. Whole-mount RNA *in situ* hybridization (WMISH) for the *Platynereis MIP* precursor (red) counterstained for acetylated tubulin (white) or DAPI nuclear stain (blue). All images shown in ventral view, with head to top. **(A)** 14 days **(B)** 14 days with ventral nerve cord region cut away to reveal digestive system **(C)** 14 days close-up foregut **(D)** 14 days close-up mid-and hindgut **(E)** 1 month **(F)** 1 month with ventral nerve cord region cut away to reveal digestive system **(G)** 1 month close-up foregut **(H)** 1 month close-up mid-and hindgut. **(I)** Schematic of *MIP* precursor expression on ventral side of 1 month *Platynereis*. **(J)** Schematic of *MIP* precursor expression on dorsal side of 1 month *Platynereis*. **(K)** Schematic overview of expression of *MIP* and digestive enzymes in the gut of 1 month *Platynereis*. White dashed lines indicate digestive system. Yellow dashed lines mark jaws. Scale bars: 50 µm. Abbreviations: fg, foregut; mg, midgut; hg, hindgut; ant, antenna; nsp, neurosecretory plexus; adc, anterior dorsal cirrus; pp, parapodia; vnc, ventral nerve cord; ch, chaetae; ac, anal cirrus; ph, pharynx.

**Additional file 2.**
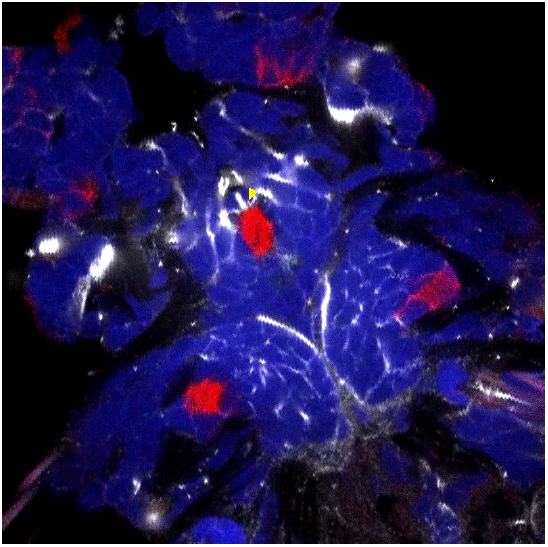
Movie of MIP expression in foregut sensory cell of 6 day *Platynereis*.mov. Whole-mount RNA *in situ* hybridization (WMISH) for the *Platynereis MIP* precursor (red) counterstained for acetylated tubulin (white) and DAPI nuclear stain (blue). Ventral view of foregut. Yellow arrowhead marks sensory dendrite of MIP-expressing cell. Green arrows indicate salivary glands.

**Additional file 3.**
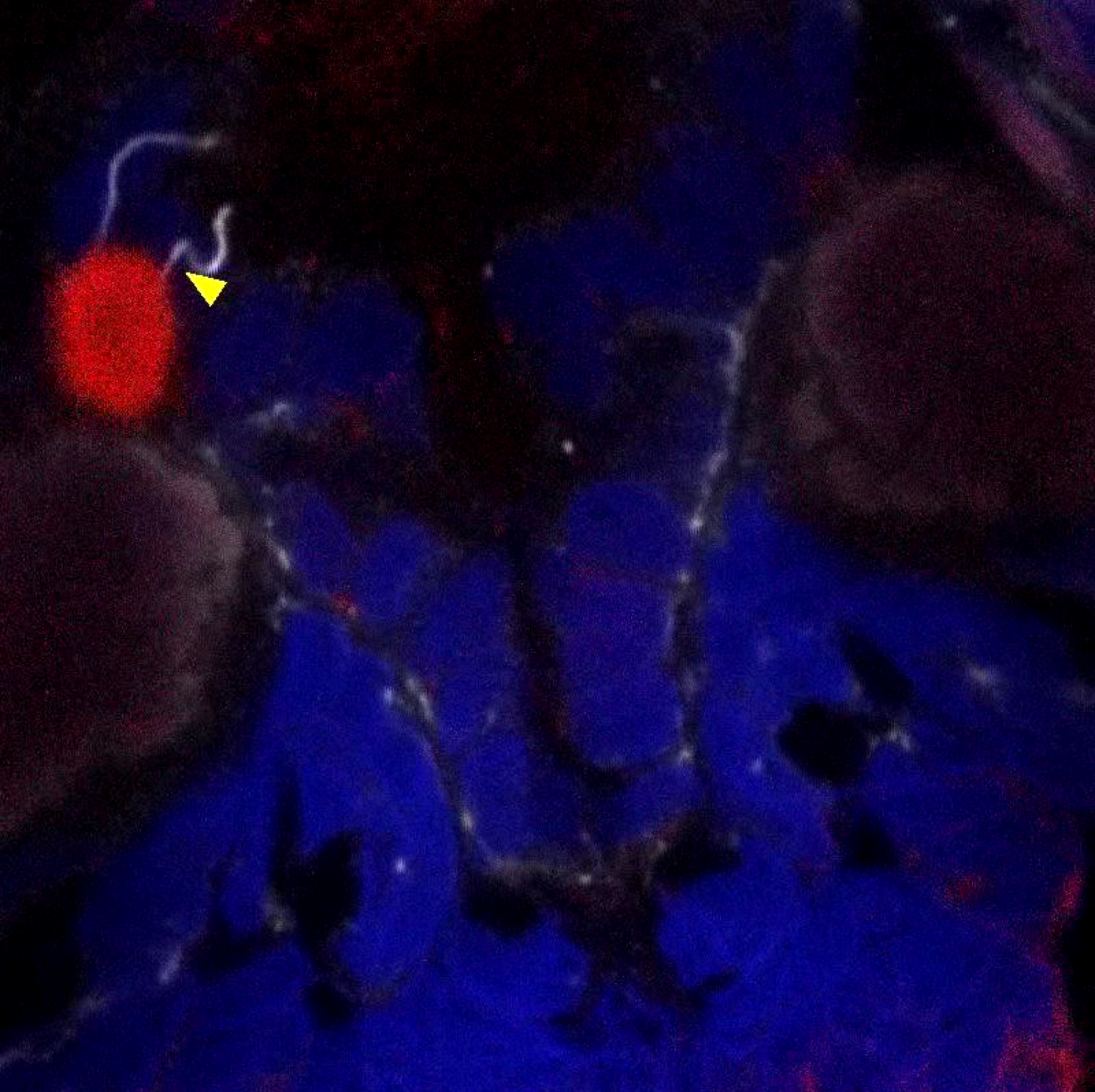
Movie of MIP expression in mid-and hindgut sensory cells of 6 day *Platynereis*.mov. Whole-mount RNA *in situ* hybridization (WMISH) for the *Platynereis MIP* precursor (red) counterstained for acetylated tubulin (white) and DAPI nuclear stain (blue). Ventral view of mid-and hindgut. Yellow arrowheads mark sensory dendrites of MIP-expressing cells.

**Additional file 4.**
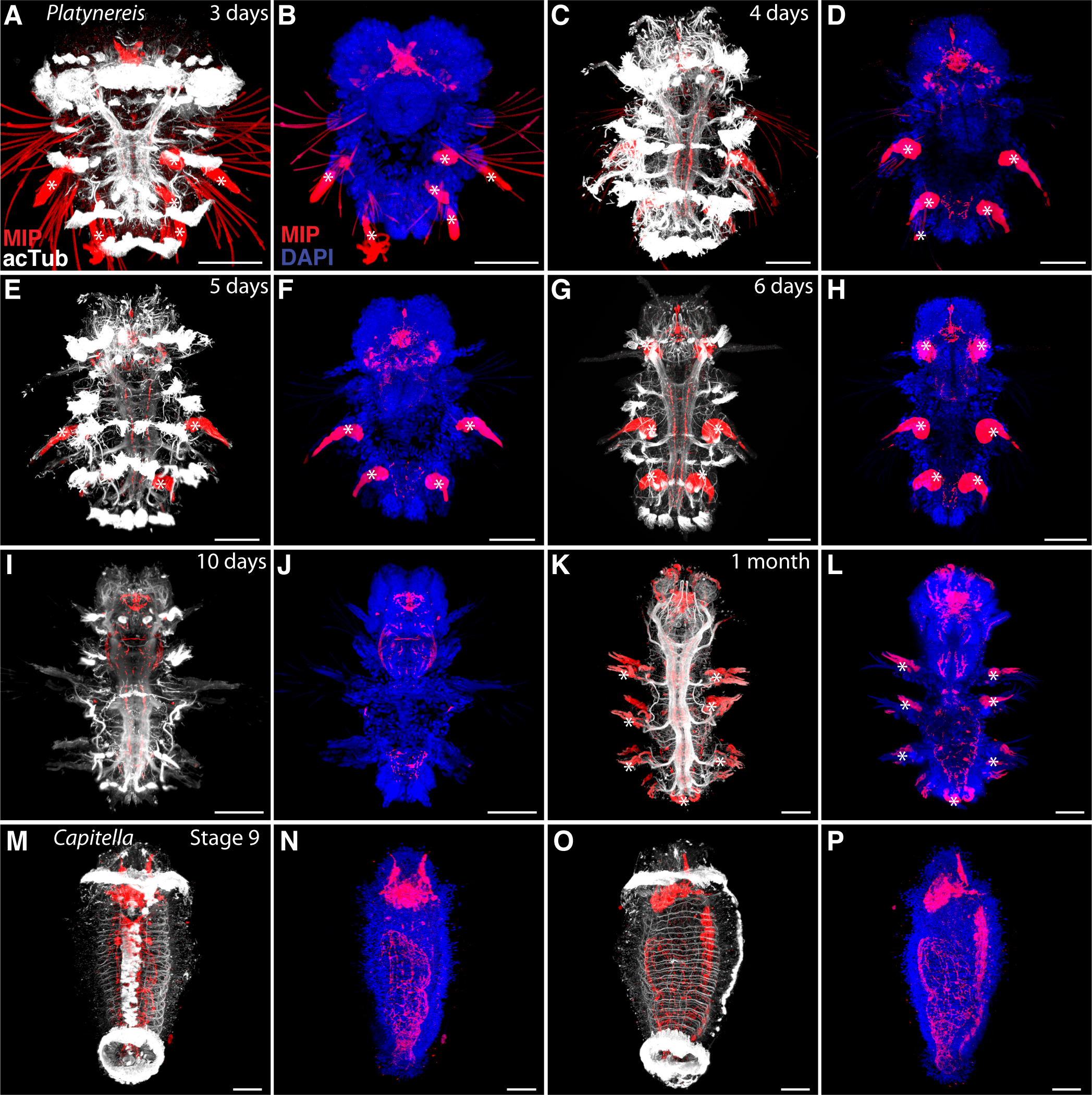
MIP peptide expression timecourse in *Platynereis*, MIP peptide expression in *Capitella* larvae.pdf. **(A-L)** Immunostaining of *Platynereis* larvae, postlarvae and juveniles with an antibody raised against *Platynereis* MIP7 (red), counterstained for acetylated tubulin (white) or DAPI nuclear stain (blue). All images ventral view with head to top. **(A)** 3 days, **(B)** 3 days with ventral nerve cord (VNC) area removed, **(C)** 4 days, **(D)** 4 days with VNC area removed, **(E)** 5 days, **(F)** 5 days with VNC area removed, **(G)** 6 days, **(H)** 6 days with VNC area removed, **(I)** 10 days, **(J)** 10 days with VNC area removed **(K)** 1 month, **(L)** 1 month with VNC removed. **(M-P)** Immunostaining of *Capitella teleta* Stage 9 larvae with an antibody raised against VWamide (red), counterstained for acetylated tubulin (white) or DAPI nuclear stain (blue). **(M)** ventral view **(N)** ventral view with VNC area removed, **(O)** lateral view with acetylated tubulin, VNC to right, **(P)** lateral view with DAPI, VNC to right. Scale bars: 50 IJm. White asterisks mark background fluorescence of parapodia, chaetae or spinning glands.

**Additional file 5.**
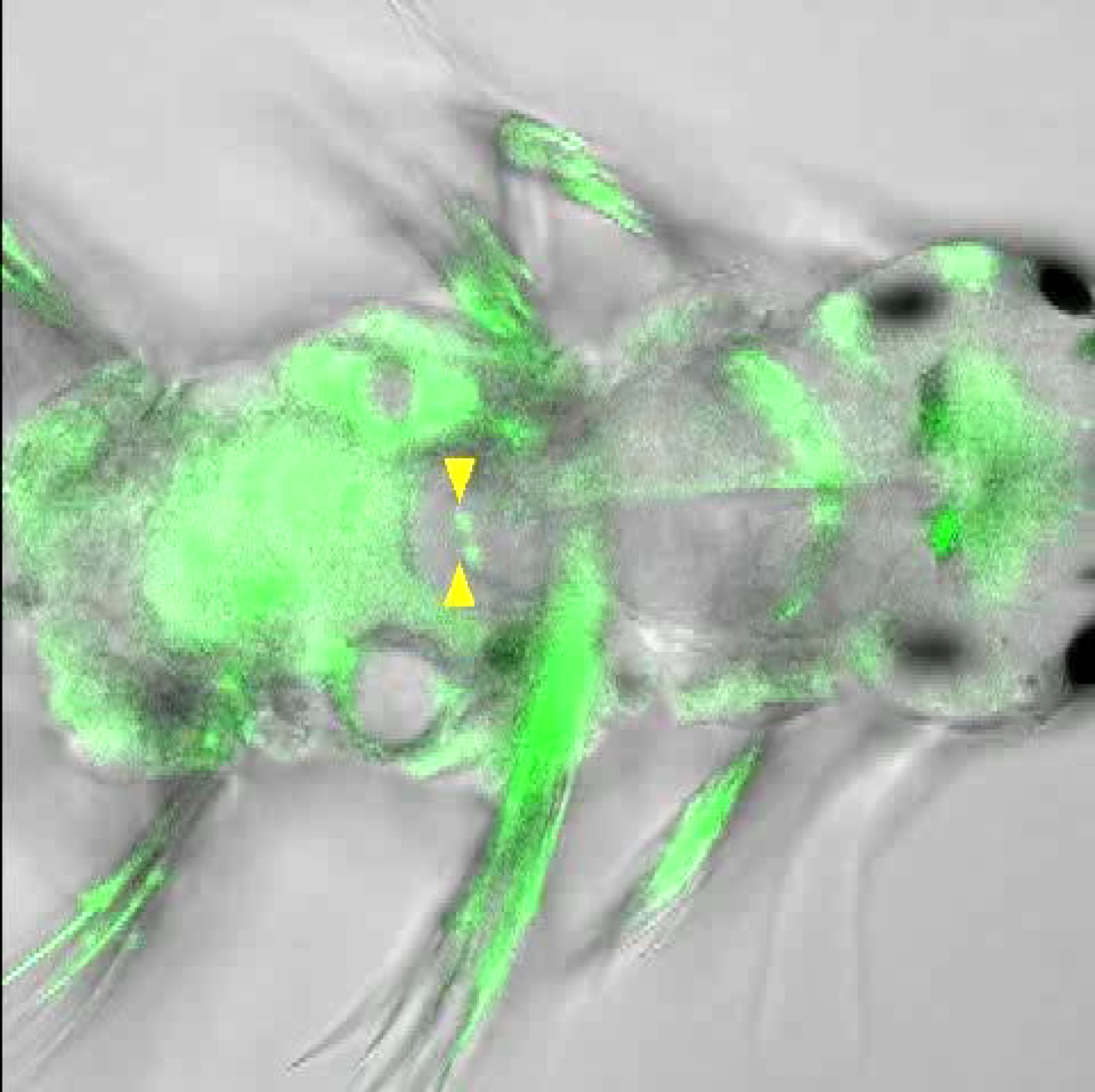
Video of *Platynereis* 6 day digestive system.mov. Calcium imaging with GCaMP6 shows muscular and neuronal activity in the digestive system of 6 day *Platynereis*. A sphincter muscle (indicated by yellow arrowheads) marks the boundary between foregut and midgut. Dorsal view with head to right.

**Additional file 6.**
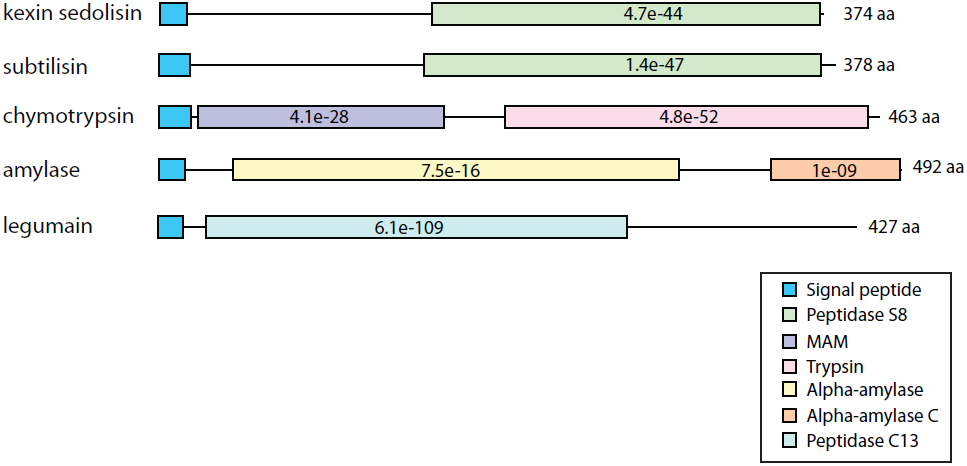
*Platynereis* digestive enzyme genes.pdf. Schematic representation of *Platynereis* digestive enzyme genes *kexin sedolisin*, *subtilisin*, *chymotrypsin*, *alpha*-*amylase* and *legumain*. Signal peptide sequence and conserved domains are marked by coloured boxes, with e-value of an HMM search against the Pfam database (http://pfam.xfam.org) indicated in box. Length of amino acid sequence obtained is shown to the right of each gene.

**Additional file 7.**
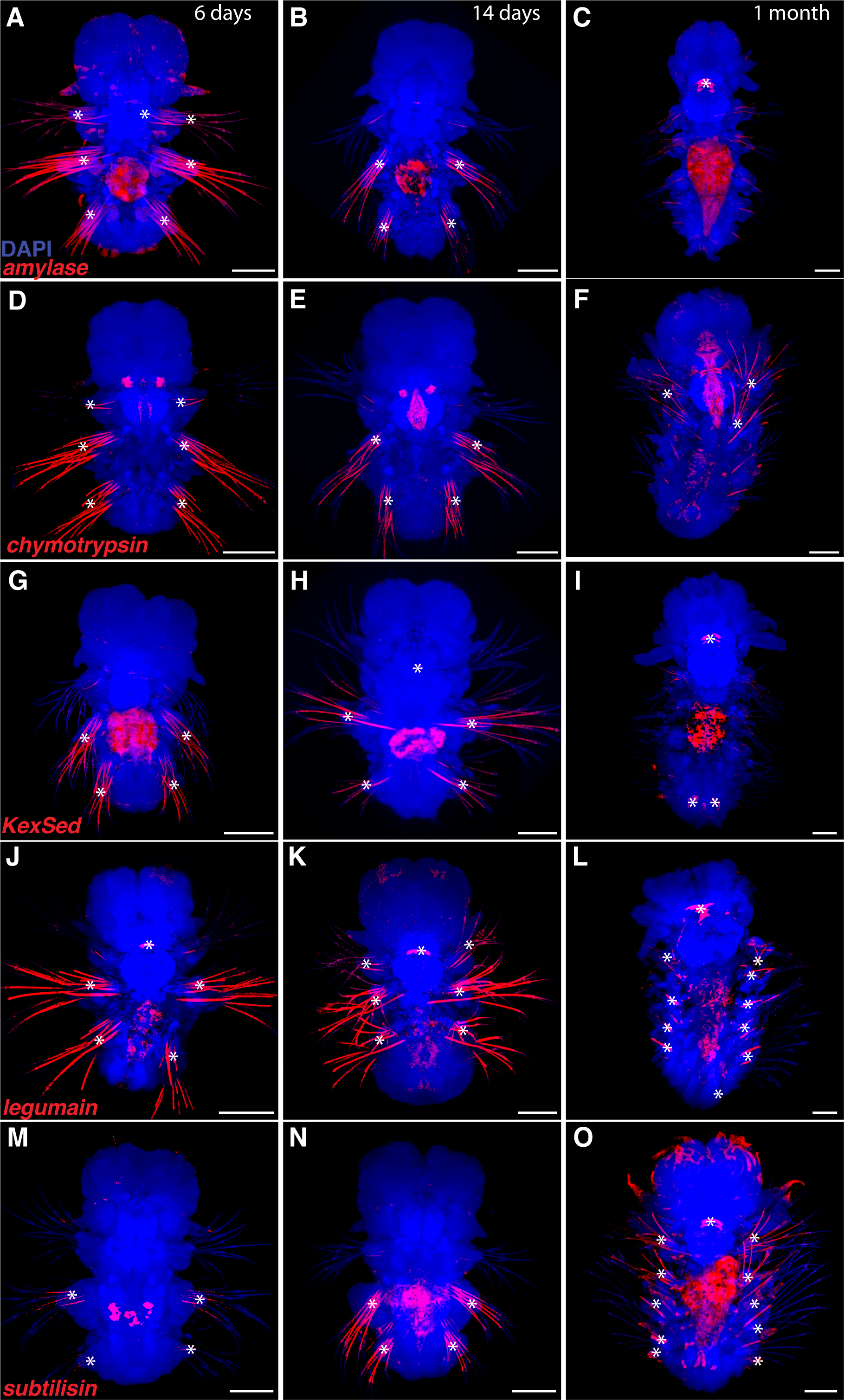
Expression timecourse of *Platynereis* digestive enzyme genes.pdf. Whole-mount RNA *in situ* hybridization (WMISH) for *Platynereis* digestive enzyme genes (red) counterstained for DAPI nuclear stain (blue). Ventral view with the head to the top, ventral nerve cord region removed to show the digestive system. (A) *alpha amylase* 6 days, (B) *alpha amylase* 14 days, (C) *alpha amylase* 1 month. (D) *chymotrypsin* 6 days, (E) *chymotrypsin* 14 days, (F) *chymotrypsin* 1 month. (G) *kexin sedolisin* (*KexSed*) 6 days, (H) *kexin sedolisin* 14 days, (I) *kexin sedolisin* 1 month. (J) *legumain* 6 days, (K) *legumain* 14 days, (L) *legumain* 1 month. (M) *subtilisin* 6 days, (N) *subtilisin* 14 days, (O) *subtilisin* 1 month. White asterisks mark background fluorescence from jaws or chaetae. Scale bars: 50 µm.

**Additional file 8.**
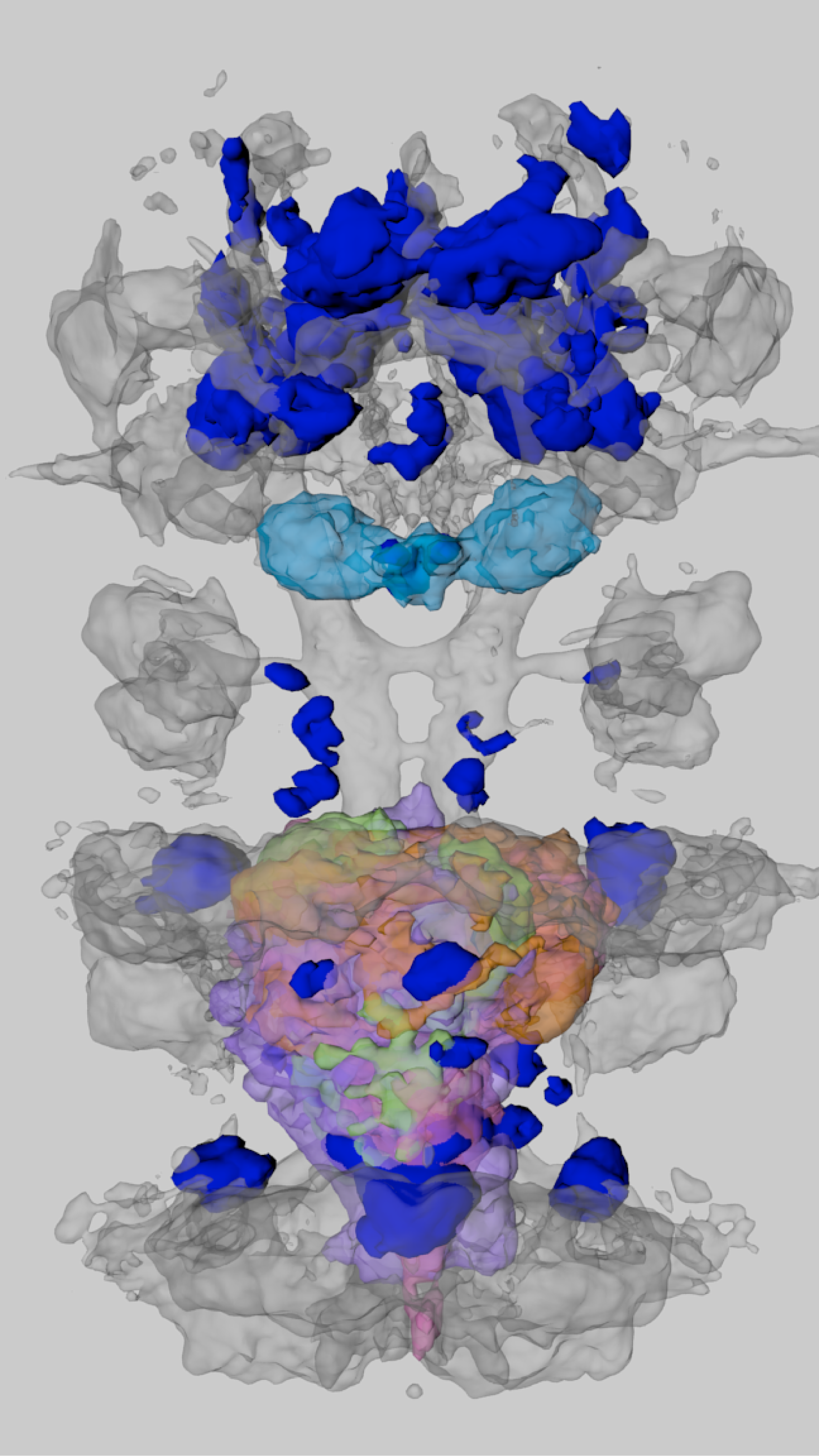
Movie of average MIP expression in 6 day *Platynereis* in relation to digestive system markers.mp4. Surface representation of the average expression domains of MIP (blue) at 6 days relative to the axonal scaffold (grey). Surface representations of digestive enzyme genes are added sequentially to mark the digestive system. In order of appearance, genes are: *alpha amylase* (orange), *chymotrypsin* (cyan), *kexin sedolisin* (green), *legumain* (purple), *subtilisin* (pink). Movie starts in dorsal view with head to the top and rotates around anterior-posterior axis.

**Additional file 9.**
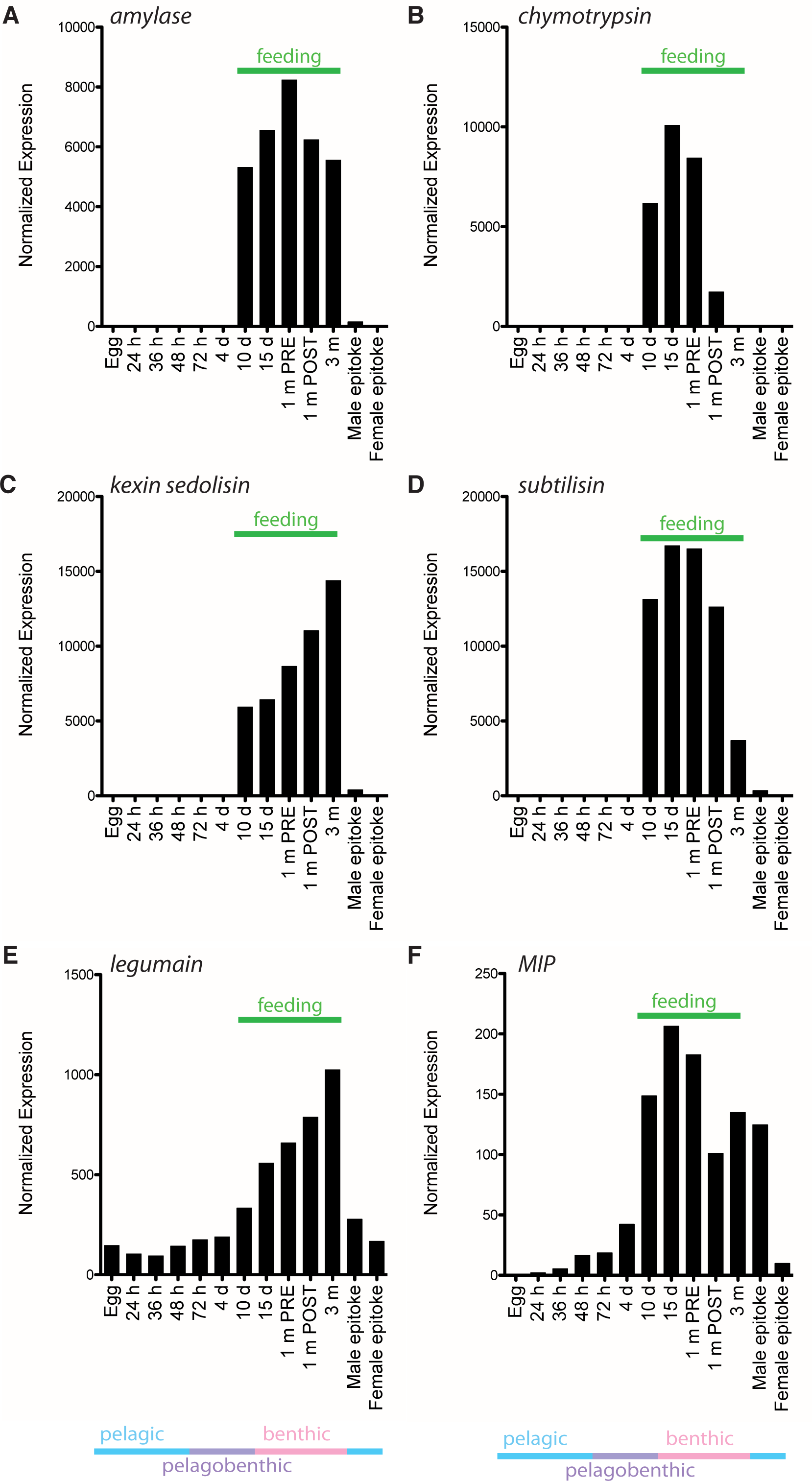
Expression of *MIP* and digestive enzyme genes throughout *Platynereis* life cycle.pdf. Histograms of normalized gene expression generated from RNA-Seq libraries of 13 different life cycle stages, from egg to sexually mature epitokes. **(A)** *alpha amylase* **(B)** *chymotrypsin*, **(C)** *kexin sedolisin*, **(D)** *subtilisin*, **(E)** *legumain* and **(F)** *MIP*. Life cycle stages during which *Platynereis* feeds are 10 days, 15 days, 1 month pre-cephalic metamorphosis, 1 month post-cephalic metamorphosis, and 3 month atokous adult. Habitat transitions are also indicated under the histograms, pelagic, free-swimming; benthic, crawling on bottom; pelagobenthic, may switch between swimming and crawling.

**Additional file 10.**
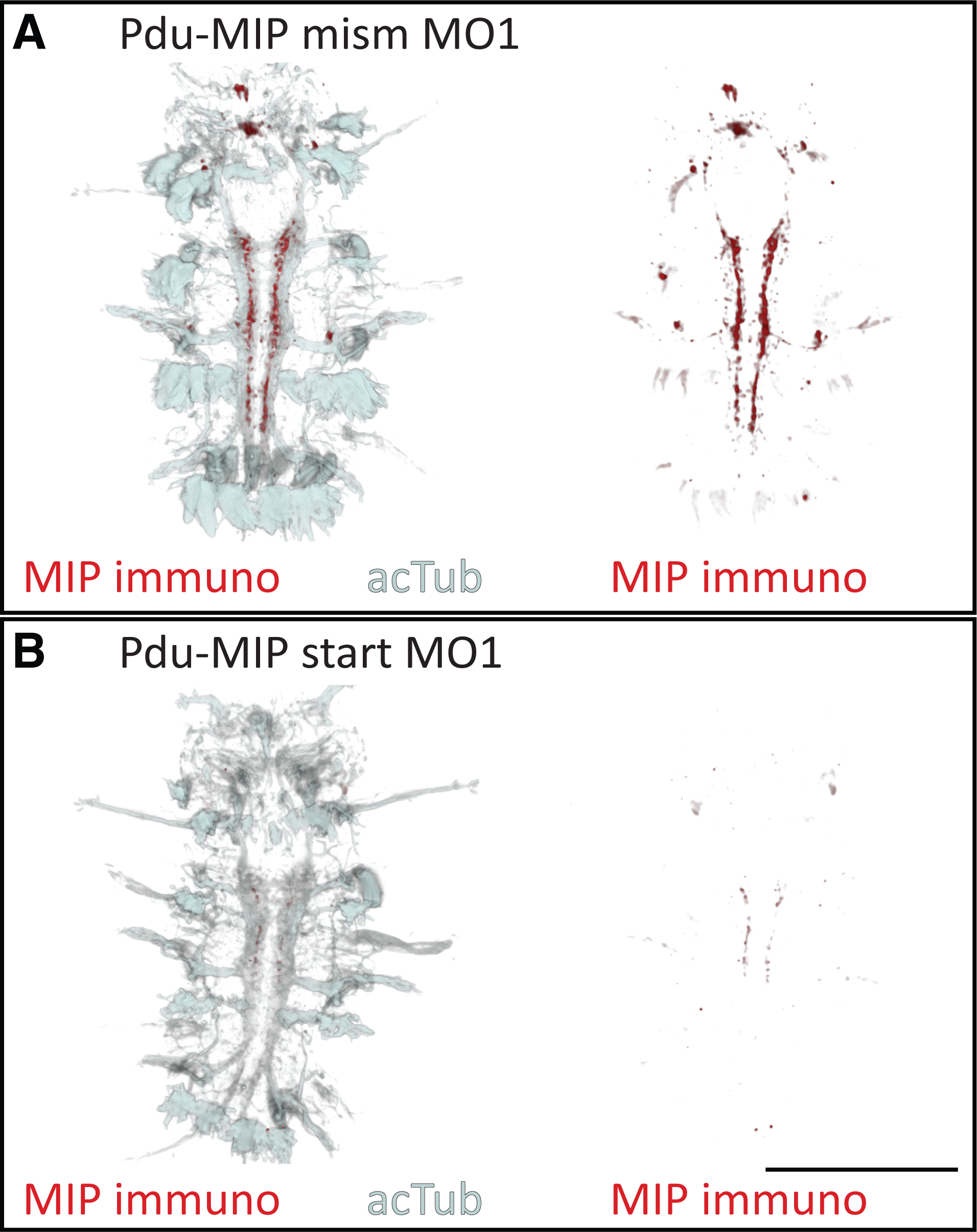
MIP knockdown *Platynereis* are morphologically similar to control *Platynereis*.pdf. (A) Histogram of length from top of head to bottom of pygidium (not including tentacular cirri) of MIP-translation-blocking morpholino-injected and MIP-mismatch-control morpholino-injected *Platynereis* at 7 – 9 days and 10 – 12 days. Data are shown as mean +/− s.e.m., n = minimum 20 larvae. (B,C) Ventral view of 6 day *Platynereis* injected with MIP mismatch or start morpholinos and immunostained with *Platynereis* MIP antibody (red) counterstained with acetylated tubulin (grey). (B) Larva injected with mismatch morpholino 1 (C) Larva injected with MIP start morpholino 1. Identical confocal microscopy and image processing parameters were applied to all images. Scale bar: 100 µm. Abbreviations: mism, mismatch; MO, morpholino; acTub, acetylated tubulin.

**Additional file 11.**
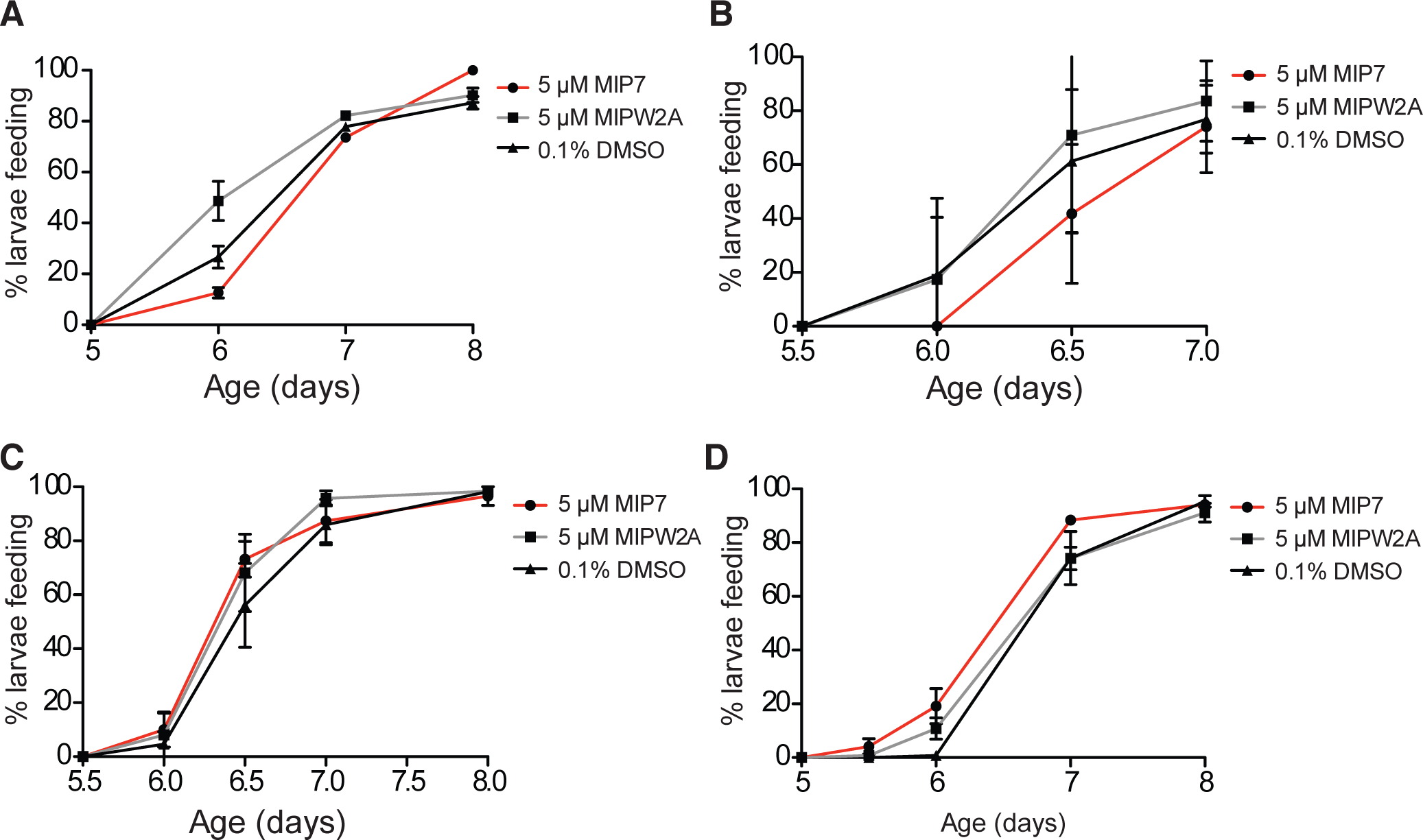
Early treatment with MIP does not induce early onset feeding.pdf. Graphs of % larvae with food in gut over time after treatment with 5 µM MIP, 5 µM control MIPW2A or 0.1% DMSO from (A) 24 hours, (B) 60 hours, (C) 4 days, or (D) 5 days. Data are shown as mean +/− s.e.m., n = 3 x 30 larvae. p-value cut-offs based on unpaired *t*-tests indicated no significant difference in initiation of feeding between MIP-treated and control larvae. MIPW2A is a control non-functional MIP peptide in which the two conserved tryptophan sites are substituted with alanines.

**Additional file 12.**
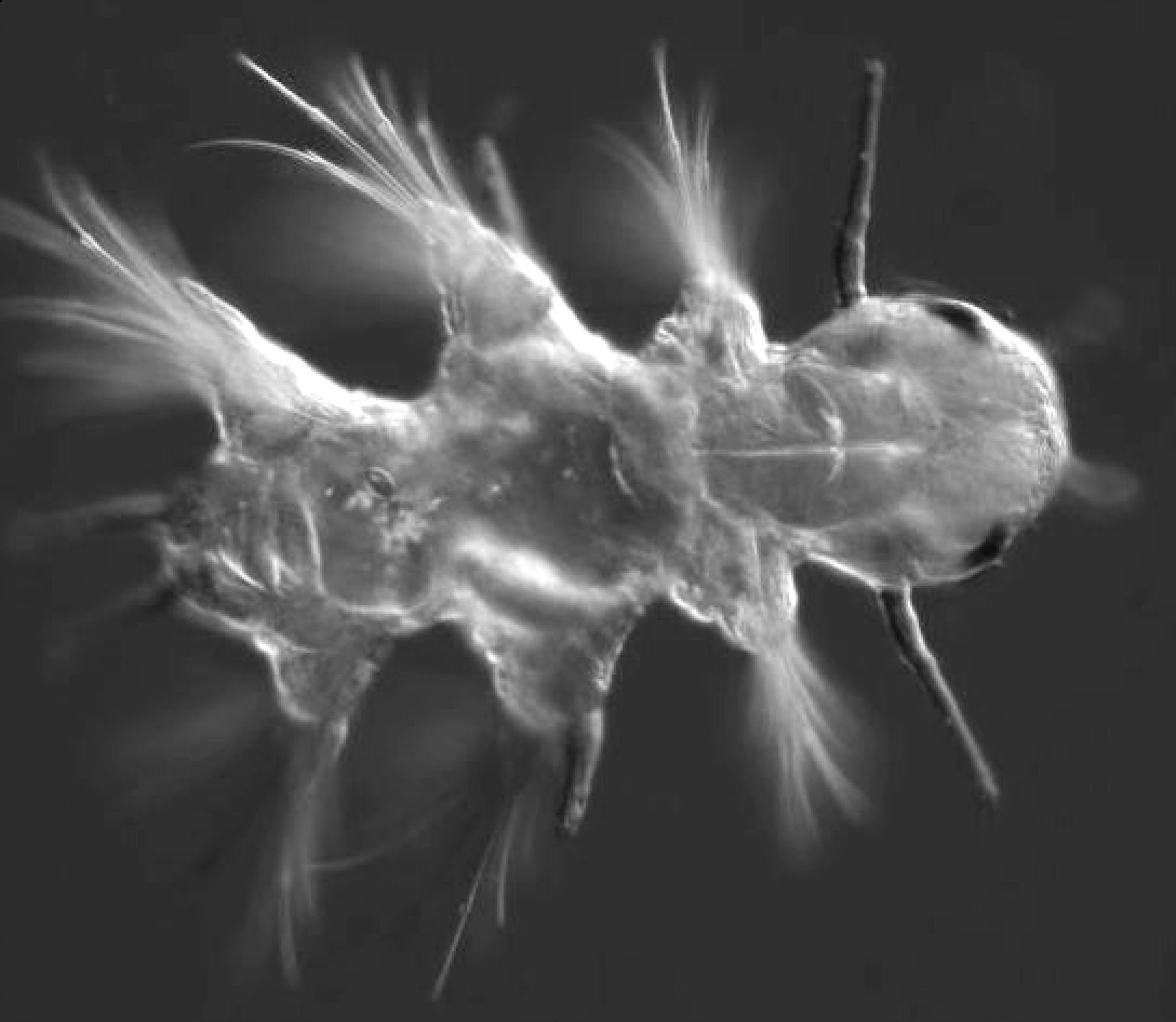
Movie of gut peristalsis in 6.5 day *Platynereis*.mov. Dorsal view with head to the right. Extension of pharynx can also be seen at 10 sec and 15 sec.

**Additional file 13.**
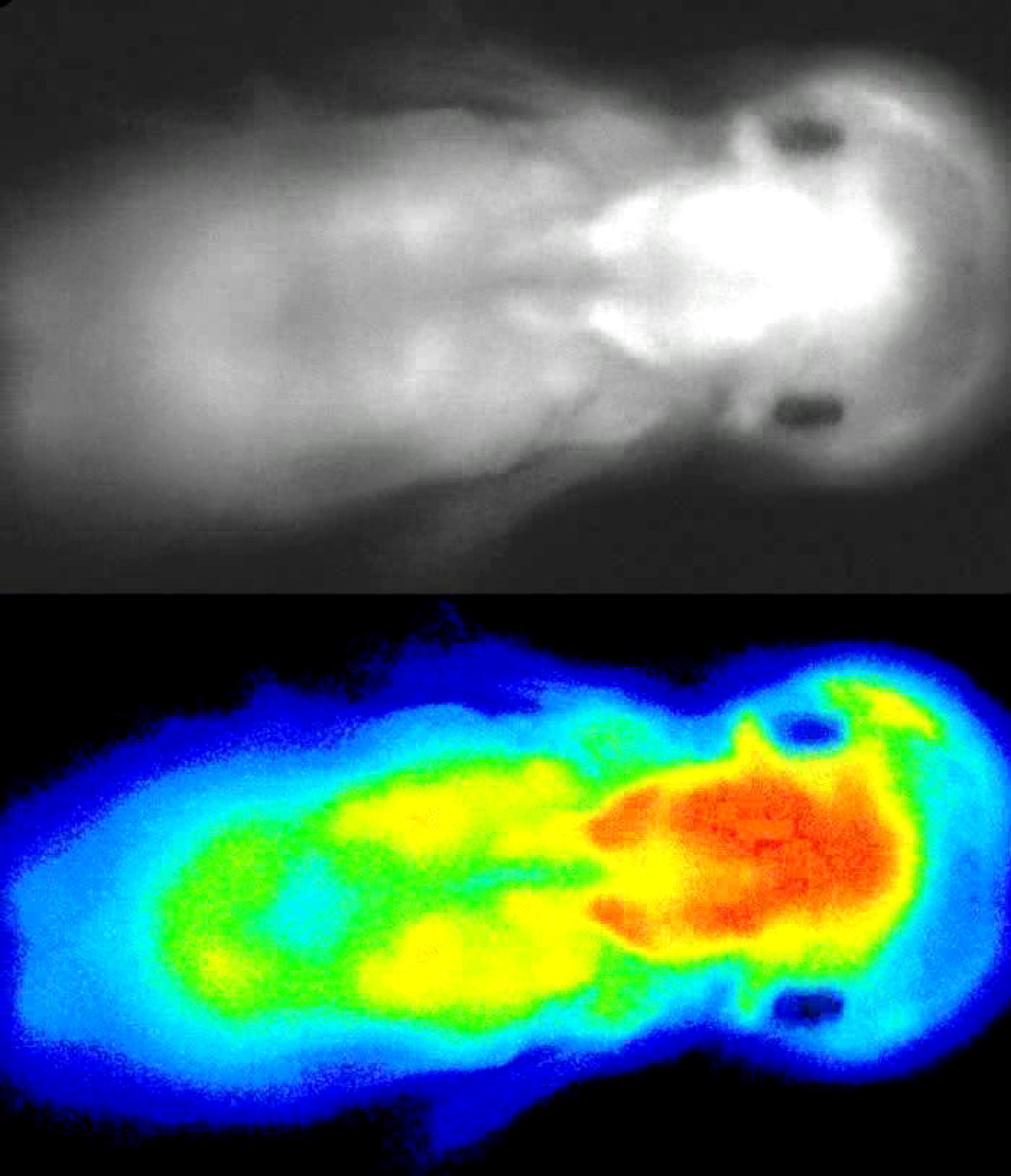
Movie of pharynx extension in 6.5 day *Platynereis*.mov. Dorsal view with head to the right. Calcium imaging with GCaMP6 shows muscular extension of the pharynx in the foregut. Top panel is differential interference contrast (DIC), bottom panel is colour-coded with Jet-LUT colour map in Image J, most intense signal in red.

**Additional file 14.**
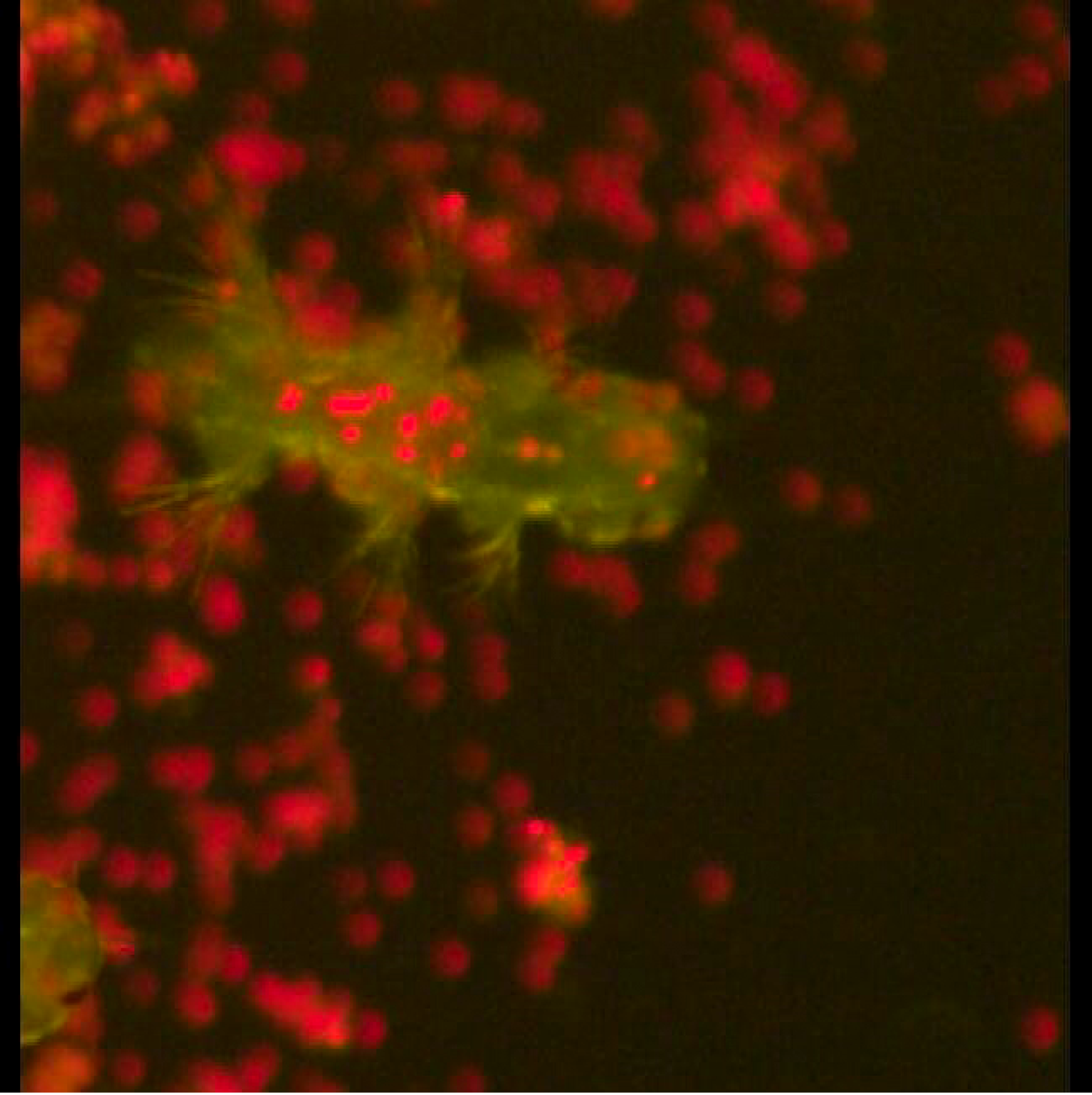
Movie of feeding in 7 day *Platynereis*.mov. *Platynereis* (green) feeds on autofluorescent *Tetraselmis* algae (red).

**Additional file 15.**
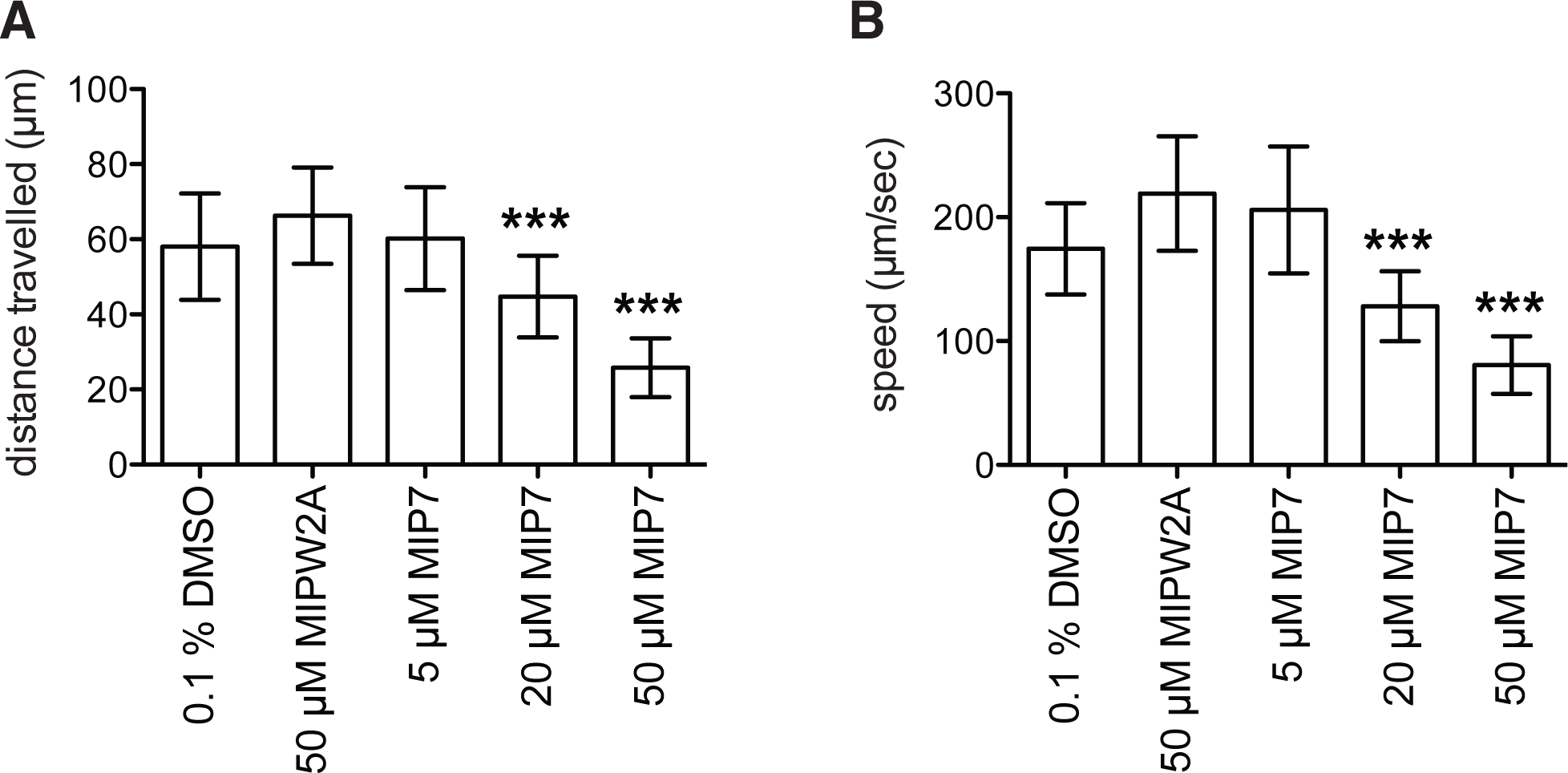
MIP treatment decreases distance traveled and speed of 6 day *Platynereis*.pdf. (A) Distance travelled by MIP-treated versus control 6.5 day *Platynereis*. (B) Speed of MIP-treated versus control 6.5 day *Platynereis*. Data are shown as mean +/− 95% confidence interval, n = 60 larvae. p-value cut-offs based on unpaired *t*-test: *** <0.001; ** < 0.01; * <0.05. MIPW2A = control non-functional MIP peptide in which the two conserved tryptophan sites are substituted with alanines.

**Additional file 16.**
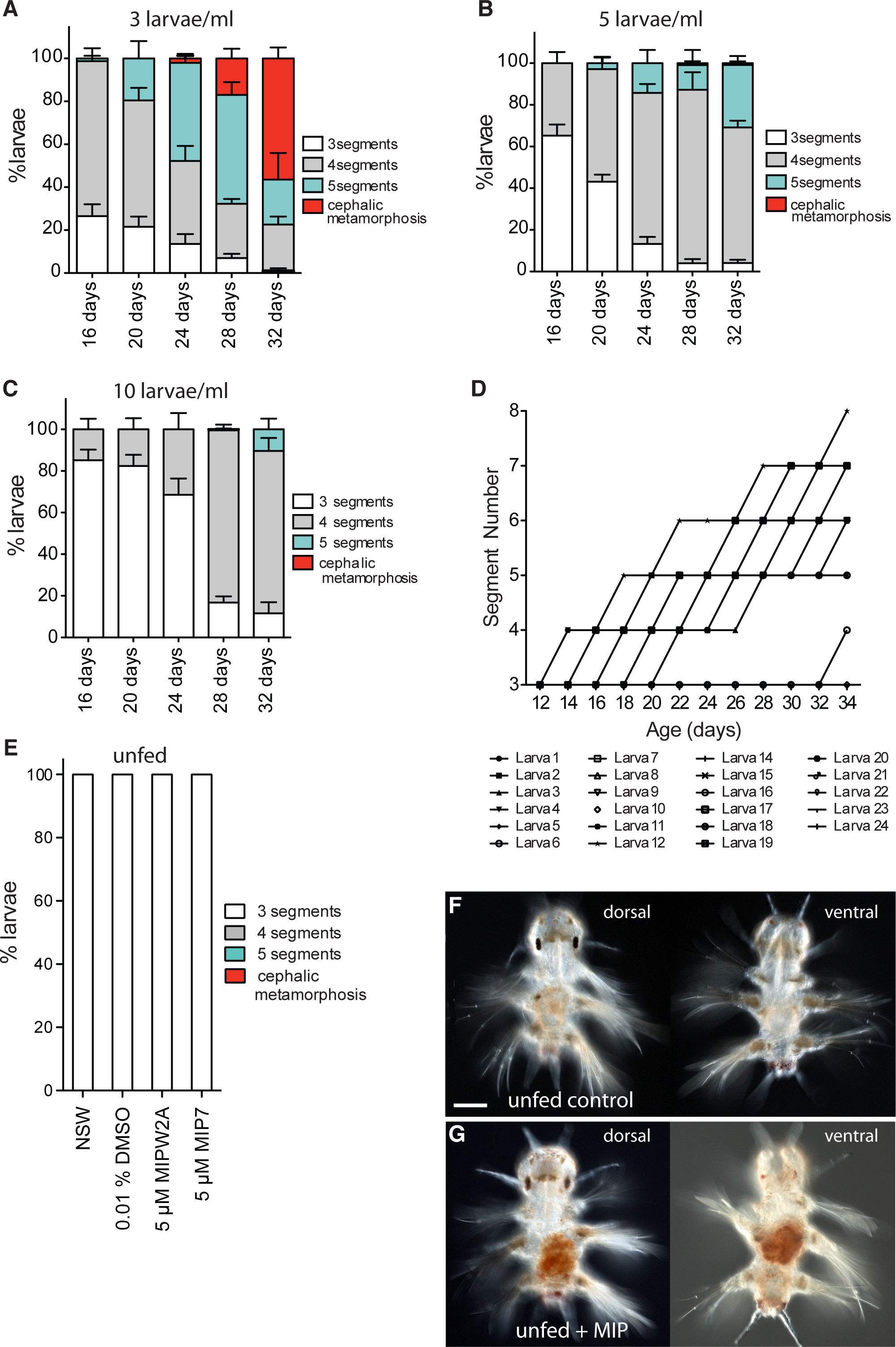
Long-term growth of *Platynereis*.pdf. Addition of posterior segments in errant juveniles raised at a density of (A) 3 larvae/mL (30 larvae in 10mL NSW) n = 3 x 30, (B) 5 larvae/mL (50 larvae in 10 mL NSW) n = 3 x 50, or (C) 10 larvae/mL (100 larvae in 10 mL NSW). n = 3 x 100. Data are shown as mean + s.e.m. All larvae were fed from 6 days. ‘Cephalic metamorphosis’ encompasses all worms which have completed cephalic metamorphosis and have 5 or more chaetigerous segments. (D) Addition of posterior segments in errant juveniles raised individually. 24 larvae were raised individually in a 24-well tissue culture dish with 2 mL NSW per well. Note: Larva #13 died prior to 12 days. (E) Addition of posterior segments in unfed 5 µM MIP7-treated and control errant juveniles. Data shown are mean +s.e.m, n = 3 x 30. NSW, filtered natural seawater. MIPW2A is a control non-functional MIP peptide in which the two conserved tryptophan sites are substituted with alanines. (F, G) Differential interference contrast (DIC) light micrographs of example unfed (F) control and (G) MIP-treated individuals at 24 days. Scale bar: 50 µm.

